# Multiomics Analysis Revealed Anti-freezing Mechanism of *Staphylococcus aureus* in Its Anti-freezing Strain

**DOI:** 10.1101/2023.10.02.560408

**Authors:** Wu Youzhi, Sun Linjun, Shi Chunlei, Jiao Ling Xia, Ye Fuzhou

## Abstract

Frozen food is currently a common food type. However, the presence of *Staphylococcus aureus* contamination caused serious challenge to frozen food safety. In this study, we explored the differences between sensitive strains and anti-freeze strains through multiomics analysis such as proteomics, phosphorylated proteomics, and metabolomics studies to understand the anti-freezing mechanism of *S. aureus*. This study compared the proteomics, phosphorylated proteomics and metabolomic differences between anti-freeze strains and sensitive strains before and after freezing. Before and after freezing, the differential protein-enriched channels changed from fructose-6-phosphate pathway, arachidonic acid metabolism pathway, atrazine degradation pathway to atrazine degradation pathway, starch and sucrose metabolism pathway, cysteine and methionine metabolism pathway, nitrogen metabolism pathway. In addition, this study inferred that *pgi* gene, *ure*A (urease subunit γ), *ure*B (urease subunit α), *ure*C (urease subunit β) gene, *mtl*D gene, *fru*B gene and *asd* gene could be crucial genes for the anti-freezing mechanism of *S. aureus*, which needs further investigation. Furthermore, the experimental results showed that PfkA, DeoC and Fda proteins from *S. aureus* were the key proteins for anti-freezing. They correspondingly involved in carbon fixation, fructose and mannose metabolism, and glycolysis/gluconeogenesis pathways in photosynthetic organisms. Finally, important metabolic pathways involved in the anti-freezing mechanism, mainly ABC transport pathway, amino acid metabolism pathway and secondary metabolite anabolic pathway.

## 1. Introduction

With the rapid development of China’s economy and the accelerated pace of daily life, people’s eating habits have begun to become simple and time-saving. In this context, quick-frozen food has become one of consumers’ favorite foods due to its convenience (Yu, Wang, Zhou, & Zhang, 2021). Quick-frozen food uses fresh agricultural products, livestock and poultry products and seafood as raw materials. After pretreatment and ingredient processing, they are quickly frozen to below −30°C with a quick-freezing device. The food can be crystallized through large ice within 30 minutes through quick-freezing. Then the core temperature drops to −18°C, and the packaged food maintains its core temperature at −18°C or below during the subsequent storage, transportation and sales process (X. J. Wang, 2013) (Miao et al., 2017). The hygienic conditions of quick-frozen food during the quick-freezing process are also strictly controlled to ensure the safety of quick-frozen food (Fang et al., 2020).

Although quick-frozen food is favored by people for its convenience, many consumers still worry about its nutritional value, health safety, and food quality. According to statistics, more than 80% of ice creams have the issues that total number of bacterial colonies exceeds the standard. Therefore, the safety and quality of quick-frozen food has become a concern among consumers. Foodborne pathogens are one of the potential safety hazards hidden in quick-frozen food. Although freezing can reduce microbial activity, there are still many microorganisms that can metabolize and cause disease under low temperature conditions. Among them, *Staphylococcus aureus* (*S. aureus*) is most likely to appear in frozen foods as contaminations. *S. aureus* infection has become a worldwide public health problem. According to European union statistics, at least 800 of the food poisoning incidents that occurred in the 1990s were caused by *S. aureus* (Wei, 2019). European countries such as Finland and Hungary report that food poisoning caused by this bacterium accounts for more than 50% of all pathogenic food poisoning; it accounts for 33% in the United States and causes at least 170,000 people poisoned every year. *S. aureus* is easily affected by physical and chemical factors such as low temperature, which can reduce the number of *S. aureus* in food and shorten its survival time in quick-frozen food (Yang, Yan, Pei, & Yang, 2020). However, *S. aureus* can often be detected in quick-frozen foods, and it has been reported that *S. aureus* can survive for up to 7 years under frozen conditions (Wagner & Byrd, 2012; S. Wu et al., 2019). Once the food is contaminated, it will remain in a bacterial contaminated state for a long time. If the bacterial contamination is not handled properly, it will easily induce diseases, and the large amount of enterotoxin produced will not disappear automatically. Therefore, it is particularly important to inhibit the growth of *S. aureus* in quick-frozen foods. In summary, the problem of *S. aureus* exceeding the standard in quick-frozen food is common and has become the main microbial hazard of quick-frozen food. It is urgent to study the anti-freeze mechanism of *S. aureus* in quick-frozen food, which can provide a theory for subsequent food processing to avoid *S. aureus* contamination.

In this study, the frozen-resistant strains and sensitive strains of *S. aureus* screened in the previous stage were used for our research. Multiomics analysis was carried out through three aspects: proteomics, phosphoproteomics and metabolomics. We compare the difference of frozen-resistant strains and sensitive strains of *S. aureus* and investigate their response to freeze condition using proteomics, phosphoproteomics and metabolomics methods. A potential anti-freezing mechanism of *S. aureus* was proposed based on our study.

## 2. Materials and methods

### 2.1 Experimental bacterial strains and culture media

The experimental strains were screened from sensitive strains no 7 and anti-freeze strains no 8 in frozen food. Tryptone Soy Broth (TSB) Medium, Tryptone Soy Agar (TSA) Medium, Brain Heart Infusion Broth (BHI), Agar, Column Waters: ACQUITY UPLC BEH Amide 1.7 µm, 2.1 mm × 100 mm column.

### 2.2 Protein extraction and digestion

The sample was extracted by SDT buffer (4% (w/v) SDS, 100 mM Tris-HCl pH7.6, 0.1 M DTT) using lysis method to extract protein. The protein was then quantified by BCA method. An appropriate amount of protein was taken from each sample for trypsin enzymolysis using the filter aided proteome preparation (FASP) method, and the peptides were desalted with C18 cartridge. After the peptides were lyophilized, 40 μL of 0.1% formic acid solution was added to redissolve them, and the peptides were quantified (OD_280_).

### 2.3 Enrichment of phosphorylated peptides

Each peptide solution was lyophilized in vacuum, and then enriched with the High-SelectTM Fe-NTA Phosphopeptides Enrichment Kit (Thermos Scientific). The enriched phosphorylated peptides were concentrated in vacuo and reconstituted with 20 μL 0.1% formic acid solution for mass spectrometry experiment.

### 2.4 LC-MS/MS data acquisition

Each sample was separated using nanoelute HPLC liquid phase system with nanoliter flow rate. buffer solution A is 0.1% formic acid aqueous solution, and buffer B is 0.1% formic acid acetonitrile aqueous solution. The chromatographic column was equilibrated with buffer A, and the sample was separated through an analytical column (homemade column, 25 cm, ID75 μm, 1.9 μm, C18) with a flow rate of 300 nL/min and a column temperature of 50 °C.

After chromatographic separation, the samples were analyzed by mass spectrometry with a timsTOF Pro mass spectrometer. The detection method is using positive ion. The ion source voltage is set to 1.5 kV. MS and MS/MS are both detected and analyzed by TOF. The mass spectrometer scan range was set at 100-1700m/z. The data acquisition mode adopts the parallel accumulation serial fragmentation (PASEF) mode. After a mass spectrometer is collected, 10 times of PASEF mode is used to collect precursor ions. The cycle window time is 1.17 seconds, and the charge number is in the range of 0-5. The spectrum, dynamic exclusion time of the tandem mass spectrometry scan was set to 24 seconds to avoid repeated scans of precursor ions.

### 2.5 Protein identification and quantitative analysis

The original data of mass spectrometry analysis was a RAW file, which was processed by MaxQuant software (version 1.5.3.17).

### 2.6 GO functional annotation

Using Blast2GO to perform GO annotation on the target protein collection, the process can be roughly summarized into four steps: sequence alignment (Blast), GO entry extraction (Mapping), GO annotation (Annotation) and InterProScan supplementary annotation (Annotation Augmentation).

### 2.7 KEGG pathway annotation and enrichment analysis

Using KAAS (KEGG Automatic Annotation Server) software to perform KEGG pathway annotation on the target protein collection.

Using Fisher’s exact test to compare the distribution of each GO classification (or KEGG pathway, or Domain) in the target protein collection and the overall protein collection and perform GO annotation on the target protein collection (or KEGG pathway, Domain) or annotation enrichment analysis.

### 2.8 Collection of metabolomics testing samples

*S. aureus* sensitive strains and anti-freezing strains, cultivated until the stationary phase, were separately inoculated onto 16-20 TSA-YE agar plates. Half of the plates were incubated overnight at 30°C in a CO_2_ incubator, while the other half were pre-frozen at −30°C to −35°C for 45 minutes and then transferred to −18°C overnight. All the TSA-YE agar plates from regular cultivation and cold stress were taken out. Approximately 400 μL of pre-chilled PBS buffer was aspirated onto each plate, and the colonies of the control strains on the TSA-YE agar plates were gently and repeatedly blown using a sterile pipette tip. The liquid from the TSA-YE agar plates was then transferred to clean 1.5 mL eppendorf tubes. The tubes were centrifuged at 12,000 rpm and 4°C for 3 minutes to collect the bacterial pellets. After discarding the supernatant, the bacterial pellets were washed at least three times with pre-chilled PBS buffer. Subsequently, the pellets were snap-frozen in liquid nitrogen for 5-10 minutes and immediately stored at −80°C in a freezer.

### 2.9 Sample preprocessing

The entire thawing process should be carried out at 4°C. Approximately 1 mL of pre-chilled methanol/acetonitrile/water solution (2:2:2 v/v) was aspirated and mixed thoroughly. The mixture was sonicated in an ice-water bath for approximately 30 minutes. It was then left to stand in a −20°C freezer for about 10 minutes, followed by centrifugation at 14,000g and 4°C for 20 minutes. During the drying process of the supernatant, a vacuum condition should be provided. For the actual mass spectrometry analysis, the sample was reconstituted in 100 μL of acetonitrile-water solution (acetonitrile:water = 1:1 v/v). After vigorous mixing and centrifugation at 14,000g and 4°C for 15 minutes, the supernatant was used for injection during the analysis.

### 2.10 Chromatography-mass spectrometry analysis

After separation using the Vanquish LC Ultra-High Performance Liquid Chromatography (UHPLC) system, the samples were analyzed using the Q Exactive series mass spectrometer (Thermo fisher Scientific) in both positive and negative ion modes using electrospray ionization (ESI). The ESI source and mass spectrometry settings were as follows: auxiliary gas heater 1 (Gas 1): 60, auxiliary gas heater 2 (Gas 2): 60, curtain gas (CUR): 30 psi, ion source temperature: 600 °C, spray voltage (ISVF): ±5500 V (positive and negative modes); the first mass-to-charge ratio detection range: 80-1200 Da, resolution: 60,000, scan accumulation time: 100 ms. The second stage employed a data-dependent acquisition method, with a scanning range of 70-1200 Da, a resolution of 30,000, scan accumulation time: 50 ms, and a dynamic exclusion time of 4 s.

### 2.11 Data processing

The raw data needs to be converted to mzX ML format. Subsequently, peak alignment, retention time correction, and peak area extraction are performed using the XCMS program. Precise mass matching (<25 ppm) and MS/MS spectral matching are used for metabolite identification, followed by searching against a custom-built laboratory database.

The data, preprocessed with Pareto-scaling, can then undergo univariate and multivariate statistical analysis. Univariate statistical analysis includes t-tests and fold change analysis, with volcano plots generated using R software. Multivariate statistical analysis includes Principal Component Analysis (PCA), Partial Least Squares Discriminant Analysis (PLS-DA), and Orthogonal Partial Least Squares Discriminant Analysis (OPLSDA). Unlike PCA, the latter two are supervised analysis methods.

### 2.12 Bioinformatics analysis

After statistical analysis, differential metabolites were identified in two comparison groups (H7 and H8, Q7 and Q8) comparison groups in both positive and negative ion modes (metabolites with VIP>1 and Fold change>1 were considered upregulated, while metabolites with VIP>1 and 0<Fold change<1 were considered downregulated; significance of the differential metabolites was assessed using a threshold of *P<*0.05). The differentially expressed metabolites were subjected to cluster analysis and KEGG pathway enrichment analysis using R software (identifying metabolite pathways with a significant difference based on a threshold of *P<*0.05).

### 2.13 Data processing and statistical analysis

Each sample was prepared with three biological replicates and three technical replicates. The significance analysis of differences between the treatment and control groups was conducted using paired t-tests in the statistical software SPSS (version 11.0). The significance level was set at *P<*0.05 or 0.01.

## 3. Results

### 3.1 Anti-freeze mechanism between sensitive and anti-freeze strains from the perspective of proteomics

There were two groups of pre-freezing samples: sensitive strains (H7) and anti-freeze strains (H8) and two groups of post-freezing samples: sensitive strains (Q7) and anti-freeze strains (Q8). Ratio analysis and one-way analysis of variance (ANOVA) were performed on data that met the criteria of having at least four out of six repeated measurements that were non-zero (*P<*0.05).

Based on the LC-ESI-MS/MS analysis results, using a threshold of *P<*0.05 and a fold change greater than 2 (upregulated > 2-fold or downregulated < 0.5) compared to the control group. For pre-freezing samples, 861 differentially expressed proteins were identified, with 423 proteins showing significant upregulation and 438 proteins showing significant downregulation. Compared to pre-freezing samples, 981 differentially expressed proteins were identified in post-freezing samples, with 543 proteins showing significant upregulation and 438 proteins showing significant downregulation.

### 3.2 GO annotation classification of differential proteins

GO enrichment analysis was performed on the differentially expressed proteins in the group of H7 and H8 before and after freezing, and the biological functions in which these proteins were involved were classified. In the category of biological processes, a higher proportion of differentially expressed proteins was associated with metabolic processes and cellular processes. In the cellular component analysis, the differentially expressed proteins were mainly related to the cell, cellular components, cell membrane, and cell membrane components. In the molecular function analysis, a higher proportion of differentially expressed proteins were involved in catalytic activity and binding activity, with 57.1% of the differentially expressed proteins belonging to catalytic activity. However, in the group of Q7 and Q8, a higher proportion of differentially expressed proteins were associated with metabolic processes, cellular processes, biological regulation, stress response regulation, biological process regulation, multi-organism processes, cellular component organization or biogenesis, localization, developmental processes, cell death, and biological adhesion in the category of biological processes. In the cellular component analysis, the differentially expressed proteins were mainly related to the cell, cellular components, cell membrane, cell membrane components, extracellular region, protein-containing complex, and organelles. In the molecular function analysis, a higher proportion of differentially expressed proteins were involved in catalytic activity, binding activity, transport activity, transport regulation activity, and antioxidant activity.

### 3.3 Enrichment analysis of differential protein pathways in sensitive strains (H7) and anti-freezing strains (H8)

In this study, KEGG annotation was performed on the differentially expressed proteins between pre-freezing sensitive strains and anti-freezing strains to analyze the main metabolic and signaling pathways in which these proteins were involved in H7 and H8, respectively. Through KEGG enrichment analysis, the top ten pathways associated with the differentially expressed proteins in pre-freezing sensitive strains. Those pathways included co-factor biosynthesis; glycine, serine, and threonine metabolism; glycolysis and gluconeogenesis; pyruvate metabolism; aminoacyl-tRNA biosynthesis; two-component system; methane metabolism; ABC transporters; fructose and mannose metabolism, and amino sugar and nucleotide sugar metabolism (Fig. 1).

**Fig. 1.**
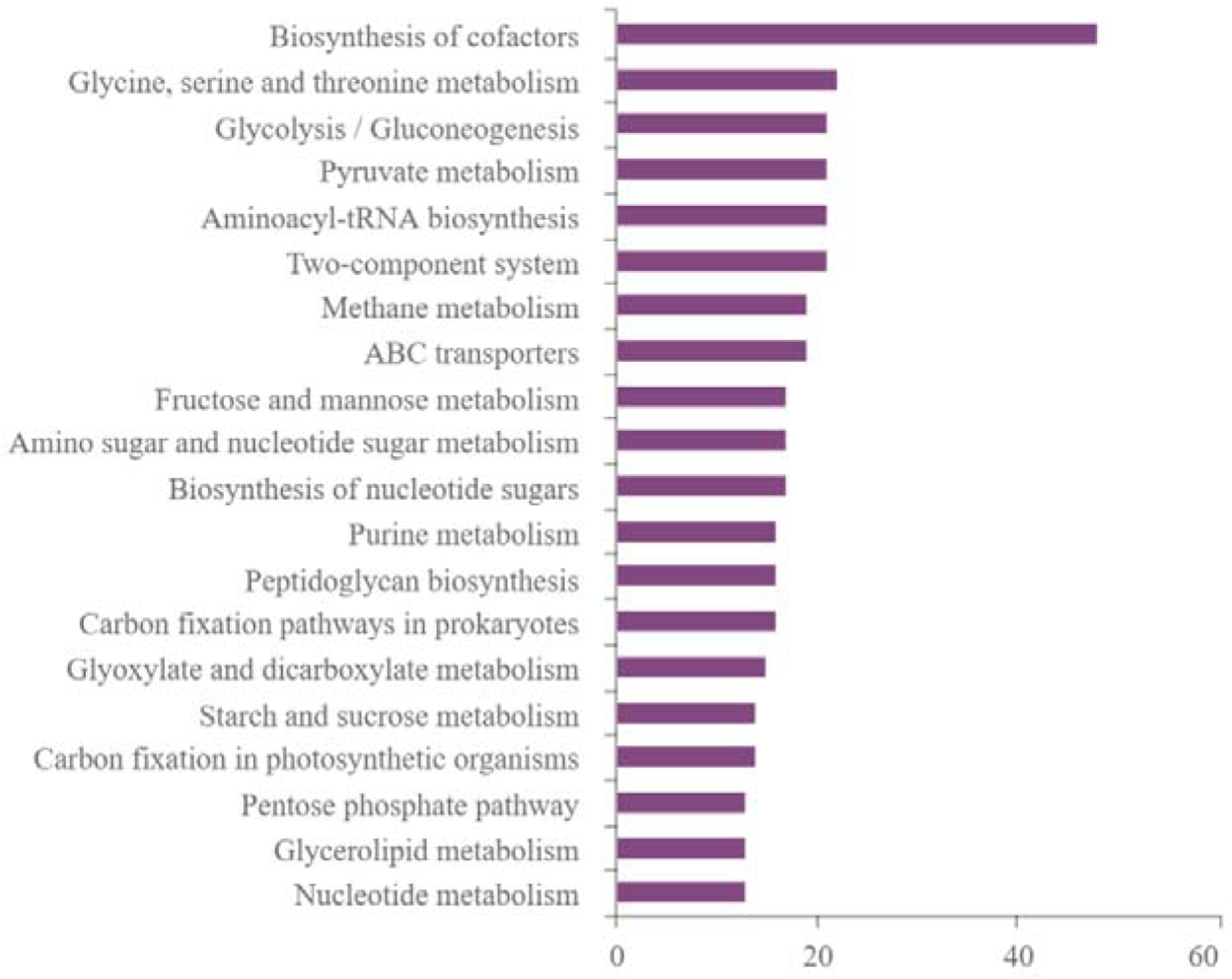
KEGG pathway annotation statistical chart of differentially expressed proteins between sensitive and antifreeze strains before freezing (Top 20 listed)

The results showed that these differentially expressed proteins were significantly enriched in the fructose-6-phosphate pathway, arachidonic acid metabolism pathway, and atrazine degradation pathway (Fig. 2). Among them, 10 differentially expressed proteins were enriched in the fructose-6-phosphate pathway, including 5 upregulated proteins, such as GAY51_04280 (mannitol-specific phosphotransferase component IIA), Fbp (fructose-1,6-bisphosphatase class 3), MW1964, SACOL2028 (putative fructokinase), HMPREF0776_1704 (mannitol-specific phosphotransferase component IIA); and 5 downregulated proteins, including T398NM01_2692 (mannose-6-phosphate isomerase), BSZ10_08650 (fructose-bisphosphate aldolase), IolC (carbohydrate kinase, fructokinase), MtlD (mannitol-1-phosphate 5-dehydrogenase), FruB (tagatose-6-phosphate kinase).

**Fig. 2.**
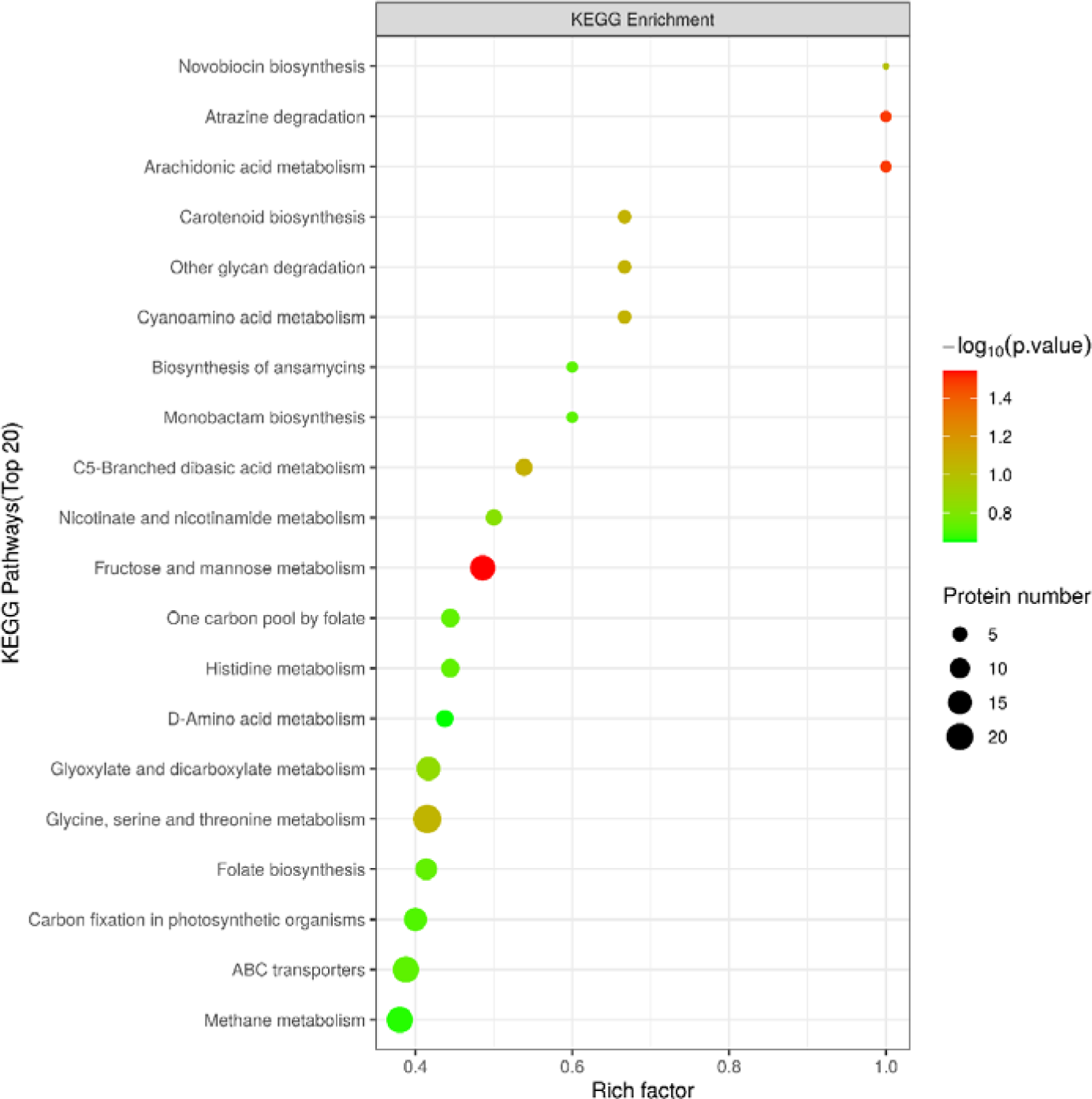
Differential expression of proteins in different biological pathway enrichment analysis between sensitive and antifreeze strains before freezing.

Two differentially expressed proteins were enriched in the arachidonic acid metabolism pathway, including One upregulated proteins, ST398NM01_2669 (glutathione peroxidase); and one downregulated protein BsaA (glutathione peroxidase). Three differentially expressed proteins, namely : ureA (urease subunit γ), ureB (urease subunit α), and ureC (urease subunit β) were enriched in the atrazine degradation pathway, all of which were upregulated

### 3.4 Enrichment analysis of differential protein signaling pathways in post-freezing sensitive strains (Q7) and anti-freezing strains (Q8)

Proteins in organisms exert their biological functions through coordinated interactions. Signaling pathway analysis helps to understand the environmental stress mechanisms of organisms comprehensively and systematically. KEGG is one of the commonly used databases for signaling pathway studies. In this study, KEGG annotation was performed on the heat stress-related differentially expressed proteins to analyze the main metabolic and signaling pathways in which these proteins were involved. Through KEGG enrichment analysis, it was found that the heat stress-related differentially expressed proteins mapped to 103 pathways, including ribosome pathways, antibiotic synthesis pathways, metabolic pathways, purine metabolism pathways, secondary metabolite biosynthesis pathways, alanine, aspartate, and glutamate metabolism pathways (Fig. 3).

**Fig. 3.**
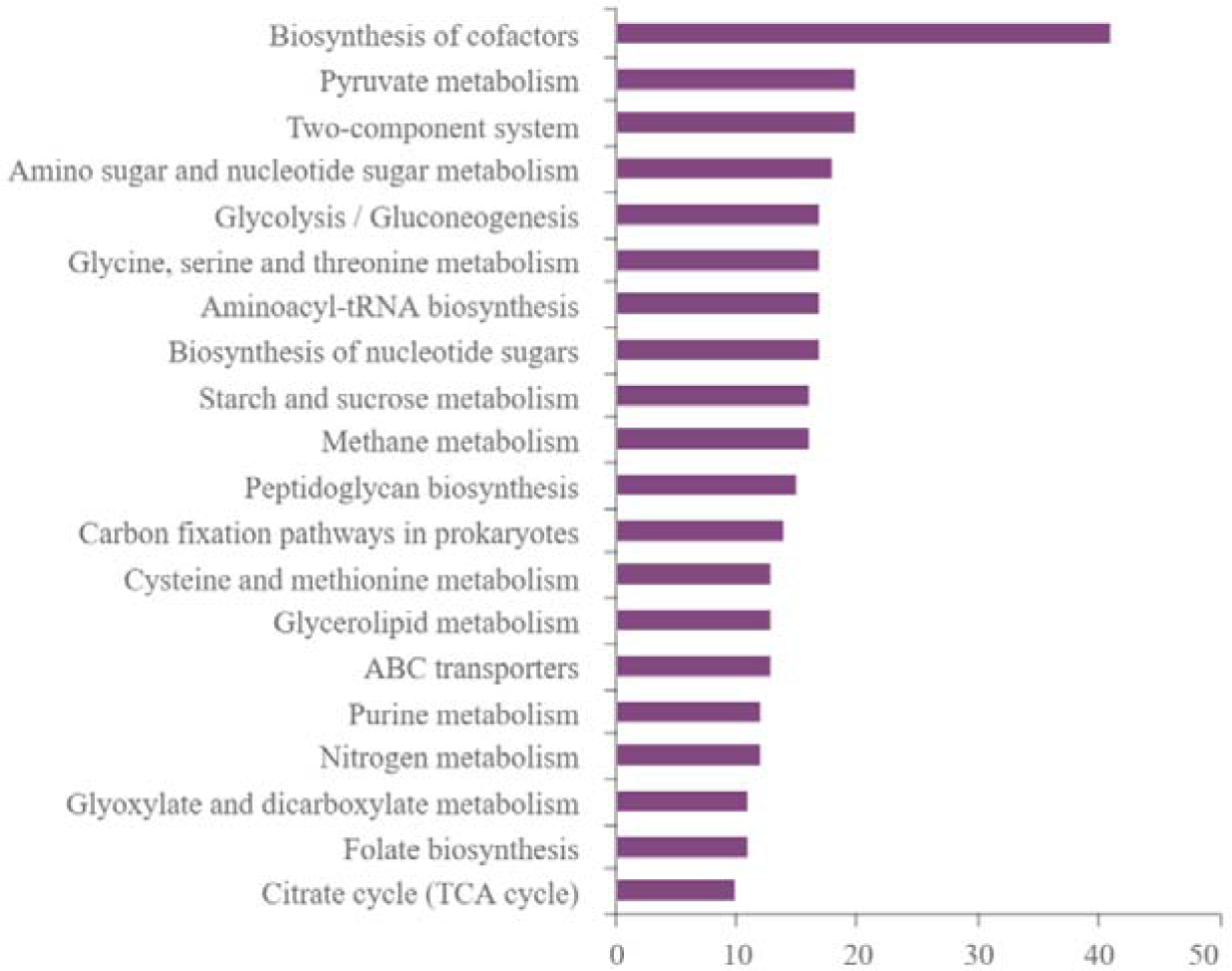
KEGG pathway annotation statistical chart of differentially expressed proteins between sensitive and antifreeze strains after freezing (Top 20)

KEGG pathway enrichment analysis was performed on the differentially expressed proteins, and the significant enrichment (*P<*0.05) of functional categories and signaling pathways of the differentially expressed proteins were demonstrated using a bubble plot. The y-axis represents functional categories or signaling pathways, while the x-axis represents ratio of the proportion of differentially expressed proteins in the corresponding functional category to the proportion of identified proteins, expressed as the log2-transformed ratio. The color of the circles represents the significance level (p-value) of enrichment, and the size of the circles represents the number of differentially expressed proteins in each functional category or signaling pathway. The results showed that these differentially expressed proteins were significantly enriched in the atrazine degradation pathway, starch and sucrose metabolism pathway, cysteine and methionine metabolism pathway, and nitrogen metabolism pathway (Fig. 4).

**Fig. 4.**
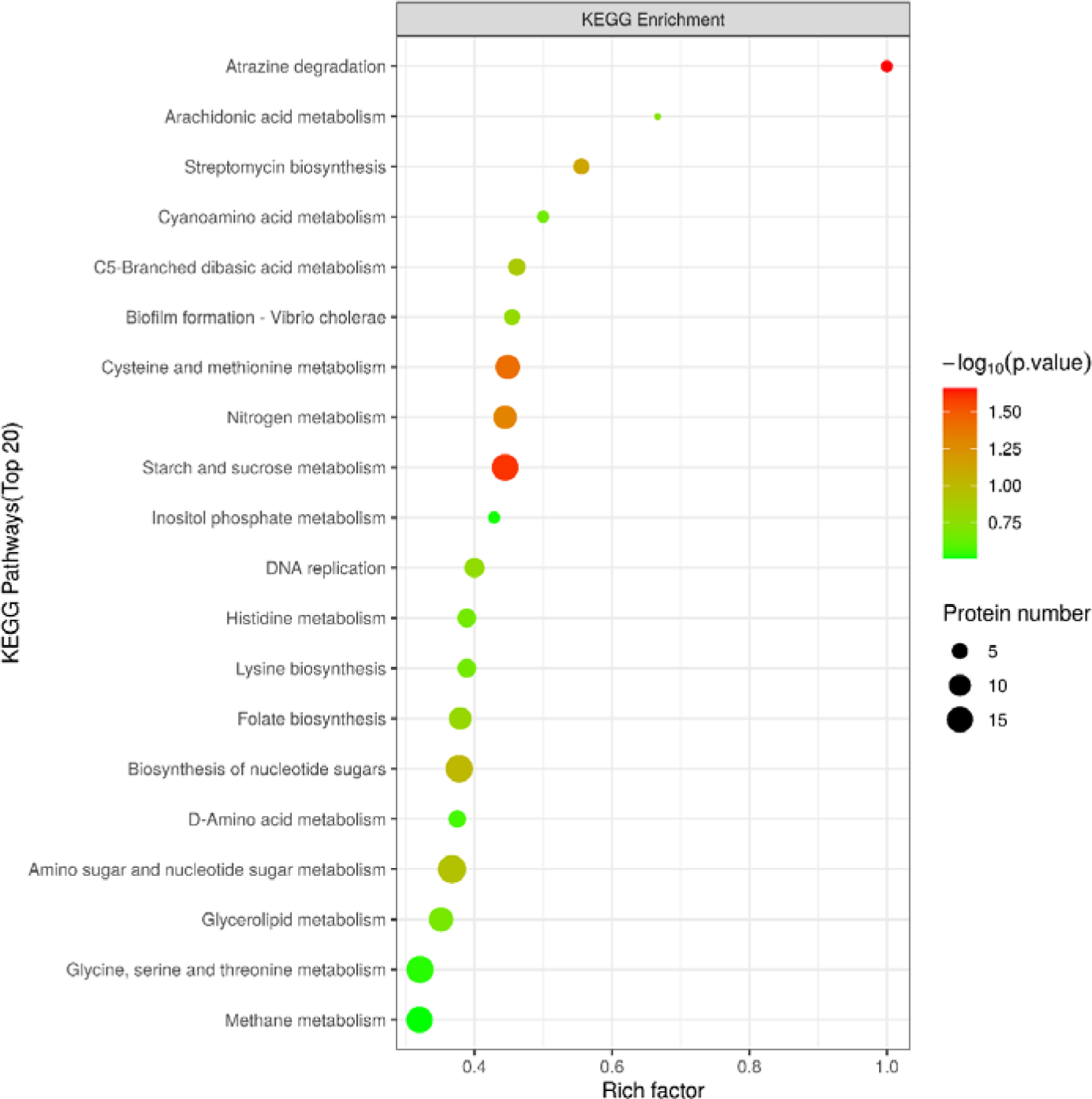
Differential expression of proteins in different biological pathway between sensitive and antifreeze strains after freezing.

Three differentially expressed proteins were enriched in the atrazine degradation pathway, all of which were upregulated, including ureA (urease subunit γ), ureB (urease subunit α), and ureC (urease subunit β).

Sixteen differentially expressed proteins were enriched in the starch and sucrose metabolism pathway, including seven upregulated proteins, such as SAB0132 (PTS system-specific EIIBC component for MurNAc-GlcNAc), DD547_01361 (cof-like hydrolase), GO941_04850 (HAD hydrolase family), MW1964 (transferase), SACOL2028 (putative fructokinase), HMPREF0776_2582 (glucokinase), Pgi (glucose-6-phosphate isomerase); and nine downregulated proteins, including ST398NM01_2577 (HAD hydrolase family), TMSFP482_19260 (sucrose-6-phosphate hydrolase), G0Y31_10190 (phosphoglucose mutase), NCTC5664_02031 (hydrolase), ScrB (sucrose-6-phosphate hydrolase), CscA (sucrose-6-phosphate hydrolase), TreA (α,α-phosphatase, trehalose-6-phosphate hydrolase), IolC (carbohydrate kinase, fructokinase).

Thirteen differentially expressed proteins were enriched in the cysteine and methionine metabolism pathway, with four upregulated proteins, including SAS1121 (guanine nucleotide phosphoribosyltransferase SAS1121), ASADH (aspartate-semialdehyde dehydrogenase), and MetB (dual function cystathionine gamma-lyase). Nine downregulated proteins included ST398NM01_2546 (pyrimidine-specific methyltransferase), MtnN (5’-methylthioadenosine/S-adenosylhomocysteine nucleosidase), BSZ10_10160 (cystathionine gamma-synthase), SerA (D-3-phosphoglycerate dehydrogenase), ST398NM01_2581 (L-serine dehydratase), ST398NM01_0629 (branched-chain amino acid aminotransferase), ST398NM01_0526 (cystathionine beta-lyase).

Twelve differentially expressed proteins were enriched in the nitrogen metabolism pathway, with four upregulated proteins, including TMSFP482_22710 (nitrate reductase large subunit), NarH (nitrate reductase). Eight downregulated proteins included ST398NM01_0277 (nitric oxide reductase subunit), NarH (nitrate reductase subunit beta), NasD (nitrite reductase), NarG (nitrate reductase), BSZ10_00700 (nitrate reductase large subunit).

### 3.5 Identification and quantification of phosphorylated proteins in pre-freezing strains (H7, H8) and post-freezing strains (Q7, Q8)

Phosphoproteomic analysis of sensitive strains and anti-freezing strains was performed. A total of 1,135 phosphorylation sites on 479 proteins were identified, of which 207 sites on 111 proteins had quantitative information in the group of H7 and H8. Using the criteria of fold change (FC) > 2.0 (upregulated > 2.0-fold or downregulated < 0.5-fold) and P value < 0.05 (T-test or other statistical tests), 66 significantly differentially phosphorylated peptides were identified, including 55 upregulated peptides and 11 downregulated peptides. These data were used for subsequent bioinformatics analysis. In the group of Q7 and Q8, a total of 1,135 phosphorylation sites on 479 proteins were identified, of which 207 sites on 111 proteins had quantitative information. Using the criteria of fold change (FC) > 2.0 (upregulated > 2.0-fold or downregulated < 0.5-fold) and P value < 0.05 (T-test or other statistical tests), 61 significantly differentially phosphorylated peptides were identified, including 39 upregulated peptides and 22 downregulated peptides. These data were used for subsequent bioinformatics analysis.

Using the criteria of fold change (FC) > 2.0 (upregulated > 2.0-fold or downregulated < 0.5-fold) and P value < 0.05 (T-test or other statistical tests), significant changes in phosphorylation levels were observed in 66 phosphorylation sites on 111 proteins between pre-freezing sensitive strains and anti-freezing strains. Among these, 55 phosphorylation sites (serine: threonine: tyrosine phosphorylation ratio of 35:16:4) showed upregulated phosphorylation levels (Table S1), while 11 phosphorylation sites (serine: threonine: tyrosine phosphorylation ratio of 6:3:2) showed downregulated phosphorylation levels (Table S2).

Proteins with significantly upregulated phosphorylation levels mainly included Fda (fructose-bisphosphate aldolase class I), GpsB (cell division cycle protein), CoaD (phosphopantothenoyl-cysteine decarboxylase), DanK (molecular chaperone protein), dagK_2 (transcription regulatory factor), NusA (transcription termination/antitermination protein), ThrS (threonyl-tRNA synthetase), RplU (50S ribosomal protein L21), EbpS (elastin-binding protein), PrfA (peptide chain release factor 1), RpsM (30S ribosomal protein S13), GapA1 (glyceraldehyde-3-phosphate dehydrogenase 1), NusG (transcription termination/antitermination protein), Pta (phosphate acetyltransferase), RpsD (30S ribosomal protein S20). Proteins with significantly downregulated phosphorylation levels included Fda (fructose-bisphosphate aldolase class I), ATP-binding domain-containing proteins, PflB (pyruvate formate-lyase), ArcA (arginine deiminase).

For post-freezing sensitive strains (Q7) and anti-freezing strains (Q8), using the criteria of fold change (FC) > 2.0 (upregulated > 2.0-fold or downregulated < 0.5-fold) and P value < 0.05 (T-test or other statistical tests), significant changes in phosphorylation levels were observed in 61 phosphorylation sites on 111 proteins between post-freezing sensitive strains and anti-freezing strains. Among these, 39 phosphorylation sites (serine: threonine: tyrosine phosphorylation ratio of 25:12:2) showed upregulated phosphorylation levels (Table S3), while 22 phosphorylation sites (serine: threonine: tyrosine phosphorylation ratio of 10:9:3) showed downregulated phosphorylation levels (Table S4).

Proteins with significantly upregulated phosphorylation levels included Fda (fructose-bisphosphate aldolase class I), D-fructose-6-phosphate aminotransferase, PflB (pyruvate formate-lyase), GpsB (cell division cycle protein), RnR (ribonuclease R), DnaK (molecular chaperone protein), RplS (50S ribosomal protein L19 fragment), ClpC (ATP-dependent *Clp* protease ATP-binding subunit), YaaA (RNA-binding protein involved in ribosome maturation), EbpS (elastin-binding protein EbpS), Rrr (ribosome recycling factor), Gnd (6-phosphogluconate dehydrogenase, decarboxylating), FtsZ (cell division protein FtsZ), RplQ (50S ribosomal protein L17). Proteins with significantly downregulated phosphorylation levels mainly included Fda (fructose-bisphosphate aldolase class I), D-fructose-6-phosphate aminotransferase, PflB (pyruvate formate-lyase), DNA-binding response regulator, and GluD (NAD-specific glutamate dehydrogenase).

### 3.6 GO annotation classification of differentially phosphorylated sites of the corresponding proteins

Gene ontology (GO) can be used to describe various attributes of genes and gene products. In this study, the distribution of differentially phosphorylated proteins in the second-level GO annotations was analyzed. For the pre-freezing sensitive strains (H7) and anti-freezing strains (H8), the results showed that in the biological process category, the differentially phosphorylated proteins were mainly involved in cellular processes, metabolic processes, biological and cellular component organization or biogenesis, regulation of biological processes, regulation of cellular processes, developmental processes, binding activity, molecular function, catalytic activity, transporter activity, and structural molecule activity (Fig. 5). For the post-freezing sensitive strains (Q7) and anti-freezing strains (Q8), The results showed that in the biological process category, the differentially phosphorylated proteins were mainly involved in cellular processes, metabolic processes, cellular component organization mainly related to cells, biological regulation and localization, regulation of biological processes, developmental processes, reproductive processes, reproduction, catalytic activity, binding activity, structural molecule activity, transporter activity, cellular component, cell, cell membrane, cell part, cellular membrane part, organelle, and protein-containing complex, among others (Fig. 6).

**Fig. 5.**
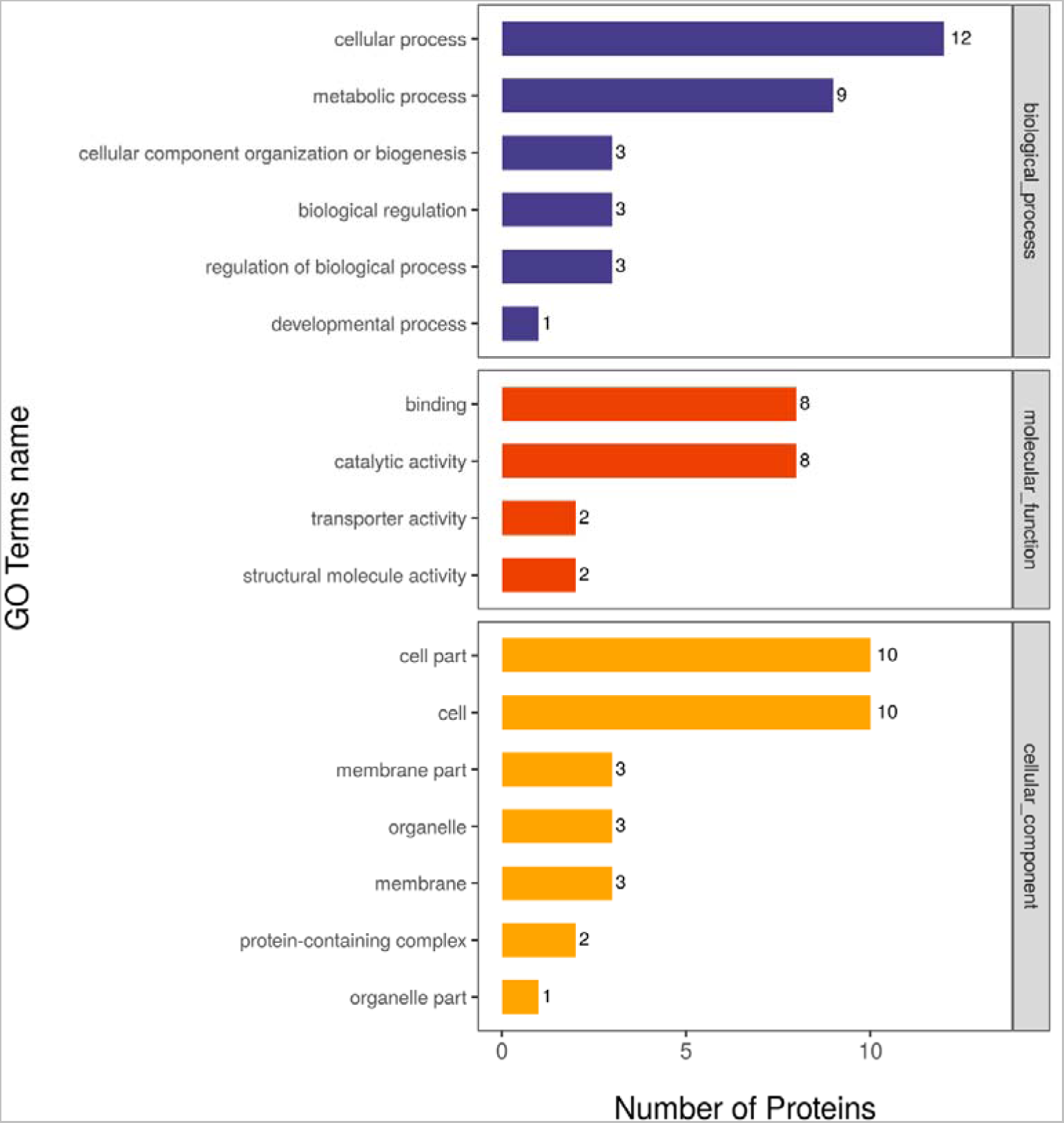
Statistical distribution of proteins corresponding to different modification sites for differential expressed proteins for sensitive and antifreeze strains before freezing in GO secondary classification.

**Fig. 6.**
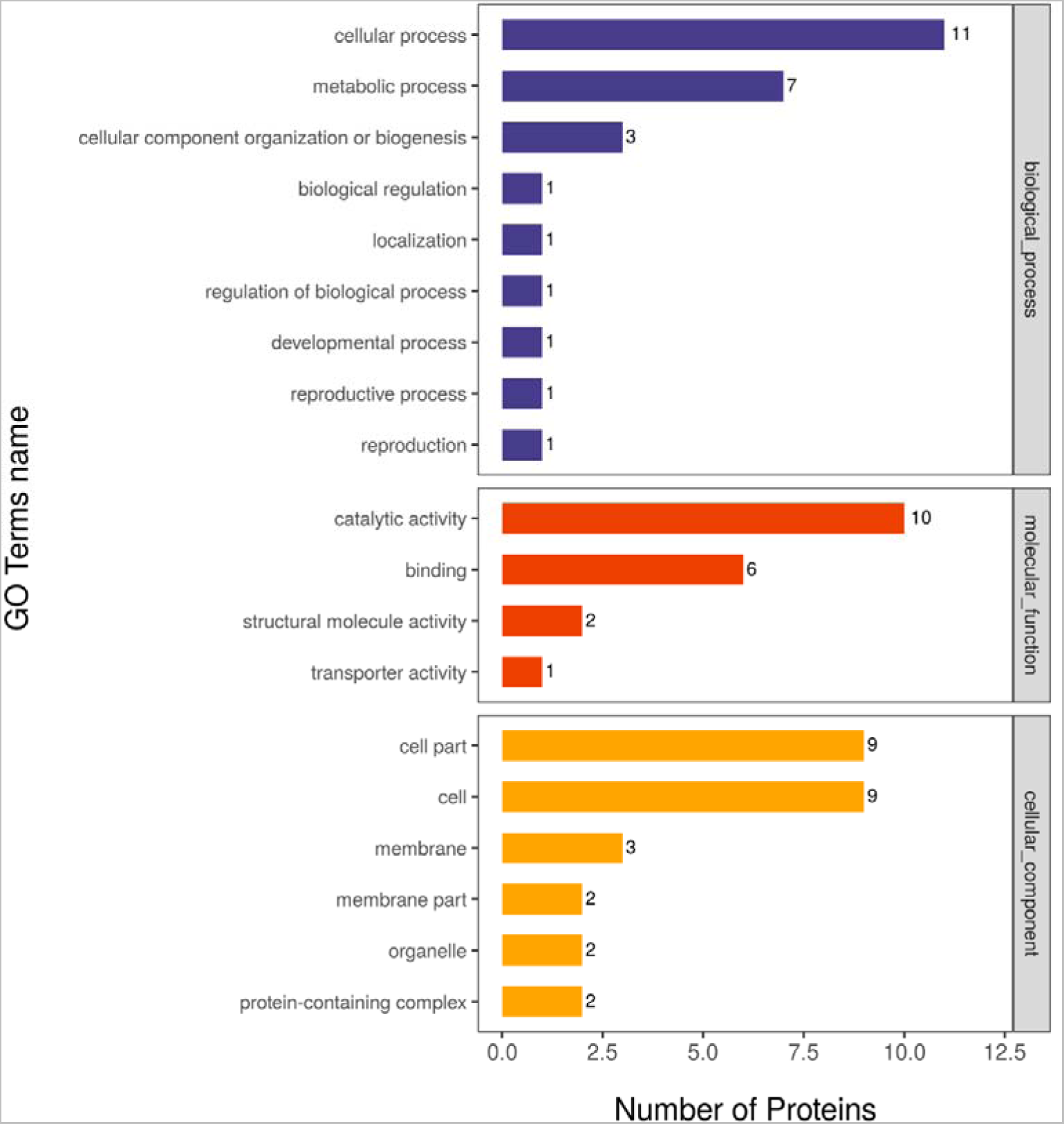
Statistical distribution of proteins corresponding to different modification sites for differential expressed proteins for sensitive and antifreeze strains after freezing in GO secondary classification.

### 3.7 KEGG enrichment analysis of differentially phosphorylated sites of the corresponding proteins

KEGG is a knowledge base for systematic analysis of gene functions, linking genomic information with higher-level functional information. In this study, the differentially phosphorylated proteins were subjected to metabolic pathway enrichment analysis. For pre-freezing sensitive strains (H7) and anti-freezing strains (H8), the phosphorylation level differentiated proteins were enriched in various pathways, including pantothenate and CoA biosynthesis pathway, glycerolipid metabolism pathway, carbon fixation pathway in photosynthetic organisms, HIF-1 signaling pathway, fructose and mannose metabolism pathway, methane metabolism pathway, phosphotransferase system pathway, two-component system pathway, butanoate metabolism pathway, glycerophospholipid metabolism pathway, taurine and hypotaurine metabolism pathway, arginine biosynthesis pathway, pyruvate metabolism pathway, pentose phosphate pathway, prokaryotic carbon fixation pathway, glycolysis/gluconeogenesis pathway, citric acid cycle pathway, ribosome pathway, coenzyme biosynthesis pathway, and aminoacyl-tRNA biosynthesis pathway. Among these pathways, Fda and GapA1 were significantly enriched in the carbon fixation pathway in photosynthetic organisms, HIF-1 signaling pathway, and glycolysis/gluconeogenesis pathway. Particularly, Fda was significantly enriched in the fructose and mannose metabolism pathway, methane metabolism pathway, and pentose phosphate pathway. Additionally, MtlF was significantly enriched in the fructose and mannose metabolism pathway, and Pta was significantly enriched in the methane metabolism pathway. PfkA, Gnd, and DeoC were significantly enriched in the pentose phosphate pathway.

In terms of post-freezing sensitive strains (Q7) and anti-freezing strains (Q8), the phosphorylation level of differentiated proteins were enriched in various pathways, including nitrogen metabolism pathway, inositol phosphate metabolism pathway, glutathione metabolism pathway, carbon fixation pathway in photosynthetic organisms, fructose and mannose metabolism pathway, pentose phosphate pathway, cell cycle pathway, two-component system pathway, butanoate metabolism pathway, taurine and hypotaurine metabolism pathway, alanine, aspartate, and glutamate metabolism pathway, arginine biosynthesis pathway, HIF-1 signaling pathway, methane metabolism pathway, RNA degradation pathway, glycolysis/gluconeogenesis pathway, pyruvate metabolism pathway, ABC transporters pathway, and citric acid cycle pathway. Among these pathways, Fda and TpiA were significantly enriched in the carbon fixation, fructose and mannose metabolism, and glycolysis/gluconeogenesis pathways, especially Fda in the pentose phosphate pathway and methane metabolism pathway. Additionally, Gnd was significantly enriched in the pentose phosphate pathway.

### 3.8 Analysis of significant differentially metabolites

First, using VIP (Variable Importance for the Projection)> 1 as the screening criterion, the differential metabolites between the two groups (H7 and H8; Q7 and Q8) were preliminarily selected. Then, univariate statistical analysis was performed to validate the significance of the differential metabolites using P < 0.05 as the criterion. The metabolites selected based on VIP > 1 and P < 0.05 were considered as significantly different metabolites, while those selected based on VIP > 1 and P < 0.1 were considered as different metabolites. For H7 and H8 group, the results showed a total of 414 significantly different metabolites, including 246 upregulated metabolites (Table S5) and 168 downregulated metabolites (Table S6). In the group of Q7 and Q8, the results showed a total of 455 significantly different metabolites, including 259 upregulated metabolites (Table S7) and 196 downregulated metabolites (Table S8).

### 3.9 KEGG enrichment analysis of differentially phosphorylated protein sites

In this study, KEGG pathway enrichment analysis was performed on significantly different metabolites to identify the enriched pathways. In the group of H7 and H8, the significantly different metabolites were enriched in various pathways, including digestive system (protein digestion and absorption pathway), amino acid metabolism (D-arginine and D-ornithine metabolism, arginine biosynthesis pathway, alanine, aspartate, and glutamate metabolism pathway, beta-alanine metabolism pathway, amino acid biosynthesis pathway, lysine degradation pathway), RNA translation (aminoacyl-tRNA biosynthesis pathway), membrane transport (ABC transporters), nucleotide metabolism (pyrimidine metabolism), carbon metabolism (pyruvate metabolism pathway, butanoate metabolism pathway), signaling pathways (mTOR signaling pathway), nervous system (lysine degradation, cAMP signaling pathway, synaptic vesicle cycle pathway, GABAergic synapse pathway), and signaling molecules and interaction (neuroactive ligand-receptor interaction pathway). In the post-freezing group (Q7 and Q8), the significantly different metabolites were enriched in digestive system (protein digestion and absorption), amino acid metabolism (alanine, aspartate, and glutamate metabolism pathway, amino acid biosynthesis pathway, arginine biosynthesis pathway, D-arginine and D-ornithine metabolism, lysine degradation pathway, glycine, serine, and threonine metabolism, glutathione metabolism pathway, amino acid biosynthesis pathway, histidine metabolism pathway, beta-alanine metabolism pathway, lysine biosynthesis pathway, tryptophan metabolism pathway, cysteine and methionine metabolism, arginine and proline metabolism), signaling molecules and interaction (neuroactive ligand-receptor interaction), RNA translation (aminoacyl-tRNA biosynthesis pathway), nervous system (GABAergic synapse pathway, synaptic vesicle cycle pathway, cholinergic synapse pathway), membrane transport (ABC transporters), nucleotide metabolism (pyrimidine metabolism), carbon metabolism (butanoate metabolism pathway), signaling pathways (mTOR signaling pathway, FoxO signaling pathway, AMPK signaling pathway, cAMP signaling pathway), vitamin and cofactor metabolism (nicotinate and nicotinamide metabolism, pantothenate and CoA biosynthesis pathway), digestive system (protein digestion and absorption, mineral absorption), sensory system (taste transduction), cell growth and death (ferroptosis pathway), energy metabolism (oxidative phosphorylation), endocrine system (renin secretion), and various biosynthesis pathways of secondary metabolites.

## 4. Discussion

The utilization of proteomics in studying sensitive and freeze-resistant strains of *S. aureus* before and after freezing provides valuable insights into the freeze resistance of *S. aureus* and the molecular mechanisms underlying freeze resistance in these strains. This study involves multiple proteins from various metabolic pathways before and after freezing, highlighting the complexity of the freezing response process.

The phosphoenolpyruvate-dependent sugar phosphotransferase system (PTS) is a major carbohydrate active transport system, where sugar substrates undergo phosphorylation catalyzed by transport proteins. It has been reported that the PTS system in *Escherichia coli* leads to carbon catabolite repression (CCR) of metabolites (Deutscher, 2006). In the fermentation process with multiple carbon sources, bacteria preferentially utilize glucose, resulting in lower yields from other carbon sources (Luo, Zhang, & Wu, 2014). Studies have indicated that the *pgi* gene plays a key role in relieving CCR. Research by Mannan et al. showed that the knockout of the *pgi* gene leads to a decrease in the transcription level of *ptsG*, significantly weakening the glucose PTS system and reducing the intracellular EIIAGlc content (Kim et al., 2015). This relieves CCR and allows the utilization of alternative carbon sources (Shiue, Brockman, & Prather, 2015). Additionally, the knockout of the *pgi* gene not only relieves catabolite repression (Shiue et al., 2015) but also directs glucose metabolism towards the pentose phosphate pathway to provide NADPH (Fong, Nanchen, Palsson, & Sauer, 2006). Therefore, it can be inferred that the *pgi* gene plays a decisive role in the operation of the PTS system. On the other hand, the *pgi* gene in soybeans is significantly upregulated under salt stress, contributing to the survival of soybeans under stress conditions. Thus, it is speculated that the *pgi* gene is one of the key genes involved in salt tolerance in soybeans. Furthermore, studies have found a negative correlation between the expression level of the *pgi* gene and starch accumulation, but the relationship between them remains unknown and requires further investigation(K. Y. Zhou, 2020). Increased protein expression of the carbohydrate transport system PTS in gram-positive bacteria facilitates substance exchange and provides necessary material conditions for the repair process of sub lethally frozen *S. aureus* (Lolkema, Kuiper, ten Hoeve-Duurkens, & Robillard, 1993). In this study, the *pgi* gene was significantly upregulated in the starch and sucrose metabolism pathways, suggesting its crucial role as a key gene in the freeze resistance mechanism. Its upregulation promotes the operation of the PTS system, accelerating substance exchange and enabling the survival of *S. aureus* under freezing conditions. However, it is currently unclear whether the *pgi* gene in *S. aureus* leads to carbon catabolite repression, and further research is needed to investigate this relationship.

Furthermore, the *ureA* (urease subunit gamma), *ureB* (urease subunit alpha), and *ureC* (urease subunit beta) genes were significantly upregulated before and after freezing. Experiments conducted by Zhou et al.(C. Zhou et al., 2019) demonstrated the importance of urease in acid tolerance of *S. aureus*, as it hydrolyzes intracellular urea to produce NH4+, consuming protons and maintaining intracellular pH stability. Based on this conclusion, it can be inferred that the upregulation of urease system-related genes in freeze-resistant strains may contribute to their enhanced tolerance and improved freeze resistance. Urease is an enzyme with important functions in organisms, and the hydrolysis of urea to ammonium nitrogen in the cytoplasm is a significant nitrogen source for plants and microorganisms (Qin, 2017). Urease is also a key factor for successful colonization of *Helicobacter pylori* in the gastric environment, as it produces a substantial amount of urease under high-acid conditions (Y. Wu, Bai, Zhong, Huang, & Gao, 2017). Studies by Sheng et al. (Sheng, 2021) revealed that *S. aureus* maintains intracellular pH homeostasis and exhibits tolerance to acetic acid, lactic acid, hydrochloric acid, and citric acid stress through the urease system. However, there is currently no research linking the urease system to the freeze resistance mechanism in S. aureus, and further exploration is needed to establish this connection.

In summary, proteomic analysis provides insights into the freeze resistance and adaptive responses of *S. aureus* strains, highlighting the complex molecular mechanisms involved. The findings suggest potential key roles of the *pgi* gene related to the PTS system and the urease system-related *ure* genes in freeze resistance, laying the foundation for further research in this field.

*mtlD* and *fruB* gene expression under stress conditions can enhance organism’s stress resistance. Zong et al. (Zong & Yang, 2010) found that *mtlD* gene expression enhances salt tolerance in peanuts. When bacteria are under nutrient-deprived starvation conditions, kinases on the cell membrane, such as histidine protein kinases or serine/threonine protein kinases, are activated in response to external changes. Activated protein kinases phosphorylate the FruB protein. Subsequently, the complex formed by fruA and fruB can bind to cis-regulatory elements of target genes to regulate the expression of development-related genes and confer tolerance (Mao, Ding, & Wang, 1999).

The *asd* gene encodes aspartate semialdehyde dehydrogenase, a key enzyme in the biosynthesis pathways of lysine, threonine, methionine, and diaminopimelic acid (DAP), which is a major component of the gram-negative bacterial cell wall. When bacteria lack DAP, they undergo lysis and death (Y. H. Zhang, 2006). In this study, the *asd* gene in *S. aureus* was upregulated. Under freezing conditions, the upregulated *asd* gene thickens the main components of the cell wall, preventing cell rupture and apoptosis, thus conferring tolerance.

Carbohydrate metabolism plays a significant role in life activities, ensuring the supply of substances and energy required for organismal activities. Glucose is the simplest monosaccharide widely present in nature and serves as an ideal carbon source for organisms. Multiple metabolic pathways have been identified in microorganisms to accomplish glucose degradation, including glycolysis (Embden-Meyerhof-Parnas pathway), Entner-Doudoroff pathway, and pentose phosphate pathway (Hu et al., 2015). Among these pathways, glycolysis is the most important pathway for microbial sugar metabolism. Almost all microorganisms undergo glycolysis to convert glucose or other polysaccharides into pyruvate, releasing ATP and reducing power. The intermediates of glycolysis also serve as precursors for the synthesis of amino acids, lipids, and other cellular components. In glycolysis, *pfkA*, encoding 6-phosphofructokinase, plays a major role by catalyzing the conversion of fructose-6-phosphate to fructose-1,6-bisphosphate, which is one of the rate-limiting steps in glycolysis (J. Y. Wang, Zhu, & Xu, 2002). The upregulation of the *pfkA* gene in this study indicates a slower glycolytic rate, and the association between this phenomenon and the freeze resistance mechanism warrants further investigation.

Research has shown that the *deoC* gene, encoding deoxyribose aldolase, may enhance biofilm regeneration by degrading extracellular DNA matrix (Han, Zhu, & Dao, 2004). Therefore, the upregulation of the *deoC* gene is speculated to enhance the tolerance of *S. aureus* and is one of the key proteins in the freeze resistance mechanism. Currently, there is a lack of literature on the impact of the *deoC* gene on the freeze resistance mechanism of *S. aureus*, and further research is needed to fill this gap. Additionally, the *fda* gene is significantly upregulated in *S. aureus* before and after freezing, but there is a lack of research literature on its role, necessitating further exploration to understand its impact on the freeze resistance mechanism.

Studying the metabolic pathway differences between sensitive and freeze-resistant strains of *S. aureus* before and after freezing using metabolomics techniques can help understand the freeze resistance mechanism of *S. aureus* and find better ways to inhibit its survival and proliferation in frozen foods.

Research has shown that amino acids can accumulate in large quantities under stress conditions and play a protective role in cells (Jozefczuk et al., 2010; H. Li, Ma, Luo, Zhang, Han, & Hu, 2012). In this study, significant changes were observed in various amino acids and related metabolites during the growth of freeze-resistant strains of *S. aureus*, accompanied by changes in multiple amino acid metabolic pathways (such as alanine, aspartate, and glutamate metabolism, arginine synthesis, lysine degradation, arginine-proline metabolism, glycine, serine, and threonine metabolism, beta-alanine metabolism, glutamine-glutamate metabolism, glutathione metabolism, etc.). After freezing, the number of pathways involved in amino acid metabolism increased. Taken together, these findings indicate that amino acid metabolism plays an important role in the freeze resistance mechanism of *S. aureus*.

Metabolites such as N-acetylglutamate and guanidine were significantly upregulated, while arginine, N-alpha-acetylornithine, and glutamine were significantly downregulated in the arginine synthesis pathway. Glutamate and glutamine can interconvert, and glutamate can be converted to ornithine through a series of metabolic processes, entering the urea cycle. Meanwhile, ornithine can be converted to guanidine in the presence of ornithine aminotransferase. Based on the results, it can be inferred that the arginine synthesis pathway is active, leading to a significant increase in guanidine. Subsequently, guanidine can be converted to arginine via pathways involving alanine, aspartate, and glutamate metabolism, and then arginine can be cleaved into citrulline and ornithine by arginase. Citrulline can enter the important tricarboxylic acid cycle for metabolism. However, the metabolism of arginine exhibits an inhibitory state.

Research has shown that arginine metabolism has a significant impact on the formation of biofilms by *S. aureus*. During biofilm formation, *S. aureus* selectively acquires arginine from the external environment (De Backer et al., 2018; Zhu, Weiss, Otto, Fey, Smeltzer, & Somerville, 2007). Furthermore, arginine is involved in various important physiological processes, including energy metabolism, protein synthesis, and stress regulation in *S. aureus* (Junker et al., 2018). In the arginine deiminase (ADI) pathway, arginine is the precursor of citrulline and can also generate ornithine. Therefore, acid-adapted *Lactobacillus delbrueckii* subsp. bulgaricus can survive under high salt (180 g/L NaCl) conditions (P. J. Li, Zhao, Peng, & Cao, 2022). This suggests that arginine allows bacteria to survive in stressful environments. It is speculated that the generation of arginine may be related to the freeze resistance mechanism of *S. aureus*, but how it affects the freeze resistance mechanism is still unclear. Additionally, arginine can also be metabolized through the pathways involving arginine and proline metabolism.

Studies have shown that increased levels of proline can “awaken” bacteria in a dormant state (W. Zhang, Yamasaki, Song, & Wood, 2019). In other words, under stressful conditions, proline can reduce the sensitivity of *S. aureus* to the external environment, thereby promoting the formation of freeze-resistant strains.

In summary, significant changes were observed in various amino acids and related metabolites in the arginine metabolism pathway, indicating the important role of this pathway in the freeze resistance mechanism of *S. aureus*, which warrants further investigation.

Adenosine triphosphate-binding cassette transporters (ABC transporters) are membrane-bound proteins containing an ATP-binding cassette. They utilize the energy released from ATP hydrolysis to facilitate the transmembrane transport of substances from the external environment into the cell (Davidson, Dassa, Orelle, & Chen, 2008; Heide & Poolman, 2002; Soni, Dubey, & Bhatnagar, 2020; ter Beek, Guskov, & Slotboom, 2014). ABC transporters play a crucial role in maintaining cell integrity and are important in cell differentiation, signal transduction, and pathogenesis(Soni et al., 2020). Lin Jieting et al. (Lin, 2021) found that the glycine betaine ABC transporter system in Halo bacillus is closely related to its salt adaptation under high salt stress conditions. This indicates the crucial role of ABC transporter systems in bacterial response to adverse environments. *S. aureus* exhibits increased sensitivity to the external environment under freezing conditions. Compared to sensitive strains, freeze-resistant strains of *S. aureus* are enriched in ABC transporter systems, primarily involving differences in amino acid metabolism. Amino acids, as important nitrogen sources in bacterial growth metabolism, also participate in energy metabolism as carbon sources. The amino acid transport proteins in ABC transporter systems promote the absorption of various amino acids by bacteria to meet the metabolic requirements for growth and virulence (Soni et al., 2020). The changes in multiple metabolites in freeze-resistant strains of *S. aureus* can trigger alterations in ABC transporter systems, potentially leading to changes in the transport of multiple amino acids across the cell membrane, which significantly affect bacterial growth and virulence expression in freeze-resistant strains.

Secondary metabolites are diverse and often have antagonistic effects on other microorganisms (Hurst & Kruse, 1972). They can also regulate gene expression (Price-Whelan, Dietrich, & Newman, 2006) and influence bacterial virulence(Vallet-Gely, Opota, Boniface, Novikov, & Lemaitre, 2010; Zhu et al., 2007). Pathways related to secondary metabolite synthesis are highly susceptible to external influences. Metabolomics analysis in this study revealed changes in differential metabolites involved in secondary metabolite synthesis and metabolism pathways in freeze-resistant strains of *S. aureus*, although the connections between them have yet to be fully understood.

In conclusion, significant changes occur in cell metabolism during the formation of L-form bacteria, including processes related to environmental information processing (membrane transport and signal transduction), amino acid metabolism, energy metabolism, nucleotide metabolism, coenzyme and vitamin metabolism, and genetic information processing. These changes enable the cells to adapt to the new internal and external environment and maintain the stability and growth of freeze-resistant strains. However, due to the complexity and diversity of metabolic pathways, further research is needed to elucidate the key metabolic pathways involved in the freeze resistance mechanism of *S. aureus*.

## Acknowledgements

This work was supported by Program for Innovative Research Team (in Science and Technology) in University of Henan Province, (Grant number 21IRTSTHN024, awarded to LJ) and National Natural Science Foundation of China, General Project, (Grant number 31972169, awarded to LC)

---

**Table S1.** Differential expressed proteins for sensitive and antifreeze strains before freezing.

| Gene Name | Protein Name | Position | Amino Acid |
| --- | --- | --- | --- |
|  | Fructose-bisphosphate aldolase |  |  |
| fda | class 1 | 22 | S |
| SA1_156476 | Universal stress protein | 47 | S |
| SA1_156476 | Universal stress protein | 48 | S |
|  | Fructose-1,6-bisphosphate |  |  |
| G0Y66_01330 | aldolase | 234 | T |
| gpsB | Cell cycle protein GpsB | 109 | T |
|  | Phosphopantetheine |  |  |
| coaD | adenyllyltransferase | 144 | S |
| dnaK | Chaperone protein DnaK | 481 | S |
| mtlF | EIIA | 118 | S |
| HMPREF0776_11 |  |  |  |
| 32 | Uncharacterized protein | 165 | S |
| HMPREF0776_04 | Pyridoxamine 5'-phosphate |  |  |
| 35 | oxidase family protein | 4 | S |
| BN1321_360006 | Putative enzyme | 63 | S |
| BSZ10_07095 | Amino acid permease | 9 | T |
| BSZ10_03495 | MFS transporter | 5 | S |
| BSZ10_03495 | MFS transporter | 4 | S |
| BSZ10_03495 | MFS transporter | 7 | S |
| BSZ10_03495 | MFS transporter | 8 | S |
|  | Septation ring formation |  |  |
| ezrA_1 | regulator EzrA | 441 | S |
|  | Transcription regulator |  |  |
|  | (Contains diacylglycerol kinase |  |  |
| dagK_2 | catalytic domain) | 160 | Y |
|  | Transcription |  |  |
|  | termination/antitermination protein |  |  |
| nusA | NusA | 2 | S |
|  | Transcription |  |  |
|  | termination/antitermination protein |  |  |
| nusA | NusA | 3 | S |
|  | Flavin reductase family protein |  |  |
| GO941_00885 | (Fragment) | 10 | T |
|  | Uncharacterized protein |  |  |
| GZ116_13800 | (Fragment) | 96 | S |
| G6Y24_00010 | Phage portal protein (Fragment) | 26 | T |
| G6Y24_00010 | Phage portal protein (Fragment) | 27 | S |
| thrS | Threonine--tRNA ligase | 30 | S |
|  | Fructose-bisphosphate aldolase |  |  |
| fda | class 1 | 238 | S |
| rplU | 50S ribosomal protein L21 | 57 | T |
| SFAG_02032 | DNA-binding response regulator | 210 | S |
|  | DNA-binding response |  |  |
| SFAG_02032 | regulator | 214 | T |
| SFAG_00008 | Uncharacterized protein | 25 | T |
|  | Helicase C-terminal domain- |  |  |
| SFAG_00586 | containing protein | 318 | Y |
| SA1_156476 | Universal stress protein | 130 | S |
| GO814_15010 | AAA family ATPase | 5 | S |
| GO814_15010 | AAA family ATPase | 3 | S |
|  | DUF1828 domain-containing |  |  |
| CV021_12290 | protein (Fragment) | 10 | S |
| SARG_01812 | Uncharacterized protein | 237 | T |
| SATG_00236 | 30S ribosomal protein S1 | 359 | S |
| SATG_01785 | Uncharacterized protein | 11 | S |
| SATG_01785 | Uncharacterized protein | 9 | Y |
| AS852_14245 | Uncharacterized protein | 85 | T |
| AS852_14245 | Uncharacterized protein | 77 | T |
| AS852_14245 | Uncharacterized protein | 78 | T |
| AS852_14245 | Uncharacterized protein | 84 | T |
| AS852_14245 | Uncharacterized protein | 88 | T |
| KQU62_05730 | YppE family protein | 61 | Y |
| KQU62_05730 | YppE family protein | 64 | S |
| ebpS | Elastin-binding protein EbpS | 281 | S |
| prfA | Peptide chain release factor 1 | 28 | S |
| rpsM | 30S ribosomal protein S13 | 40 | S |
|  | Glyceraldehyde-3-phosphate |  |  |
| gapA1 | dehydrogenase 1 | 210 | S |
|  | Transcription |  |  |
|  | termination/antitermination protein |  |  |
| nusG | NusG | 101 | S |
| pta | Phosphate acetyltransferase | 129 | T |
|  | Staphylococcal accessory |  |  |
| SAKOR_00663 | regulator-like protein (SarX) | 91 | S |
| SAKOR_01000 | Uncharacterized protein | 42 | T |
| rpsT | 30S ribosomal protein S20 | 63 | S |

**Table S2.** Differential proteins for sensitive and antifreeze strains before freezing (downregulated)

| Gene Name | Protein Name | Position | Amino Acid |
| --- | --- | --- | --- |
|  | Fructose-bisphosphate aldolase |  |  |
| fda | class 1 | 238 | S |
|  | ATP-binding cassette domain- |  |  |
| GO941_12265 | containing protein (Fragment) | 10 | S |
| GO941_12265 | ATP-binding cassette domain- containing protein (Fragment) | 5 | S |
|  | ATP-binding cassette domain- |  |  |
| GO941_12265 | containing protein (Fragment) | 2 | T |
| pflB | Formate acetyltransferase | 650 | S |
| pflB | Formate acetyltransferase | 666 | T |
|  | Fructose-bisphosphate aldolase |  |  |
| fda | class 1 | 292 | S |
| arcA | Arginine deiminase | 114 | S |
| / | Uncharacterized protein | 99 | Y |
| / | Uncharacterized protein | 97 | Y |
| / | Uncharacterized protein | 96 | T |

**Table S3.**
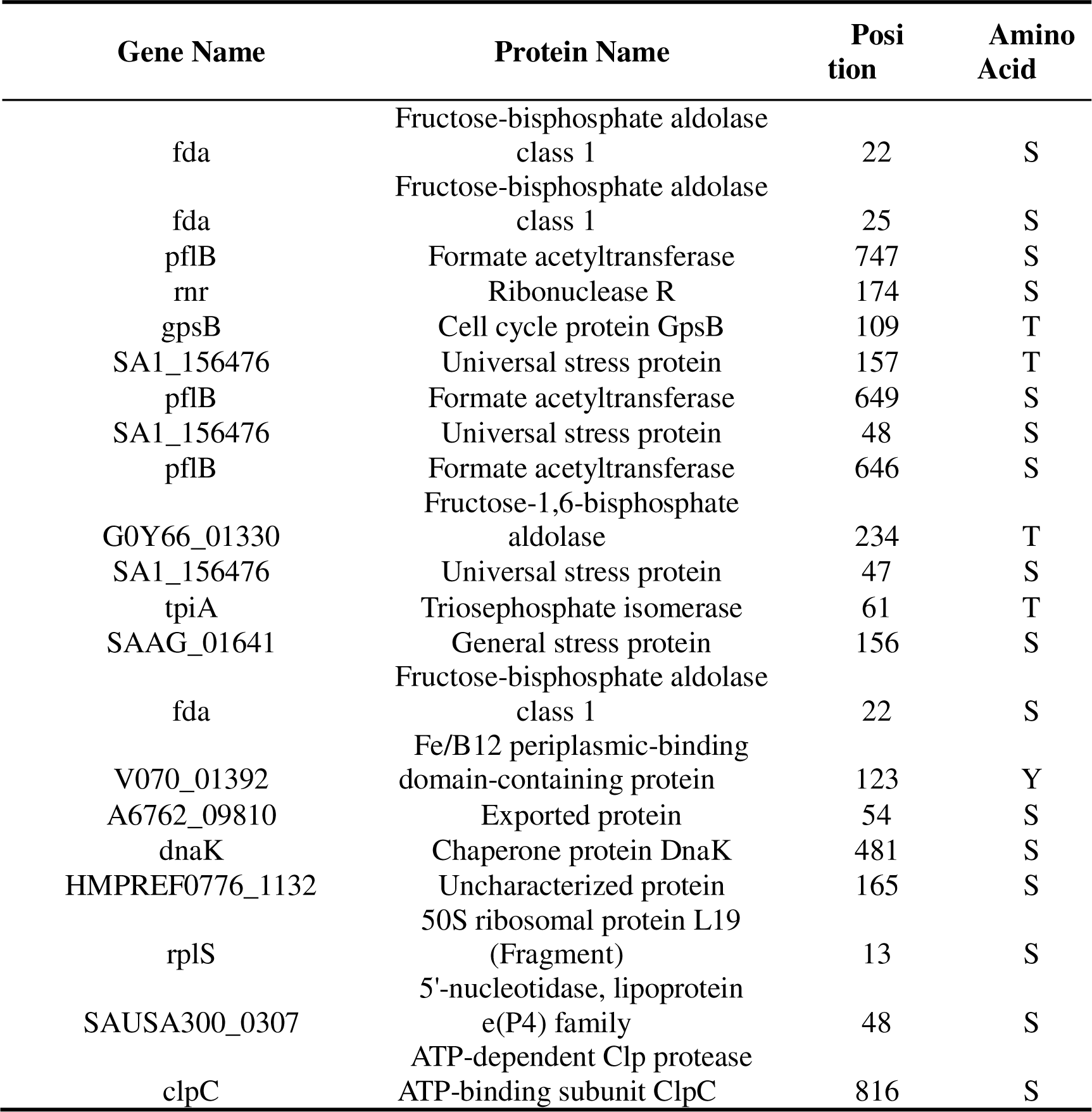
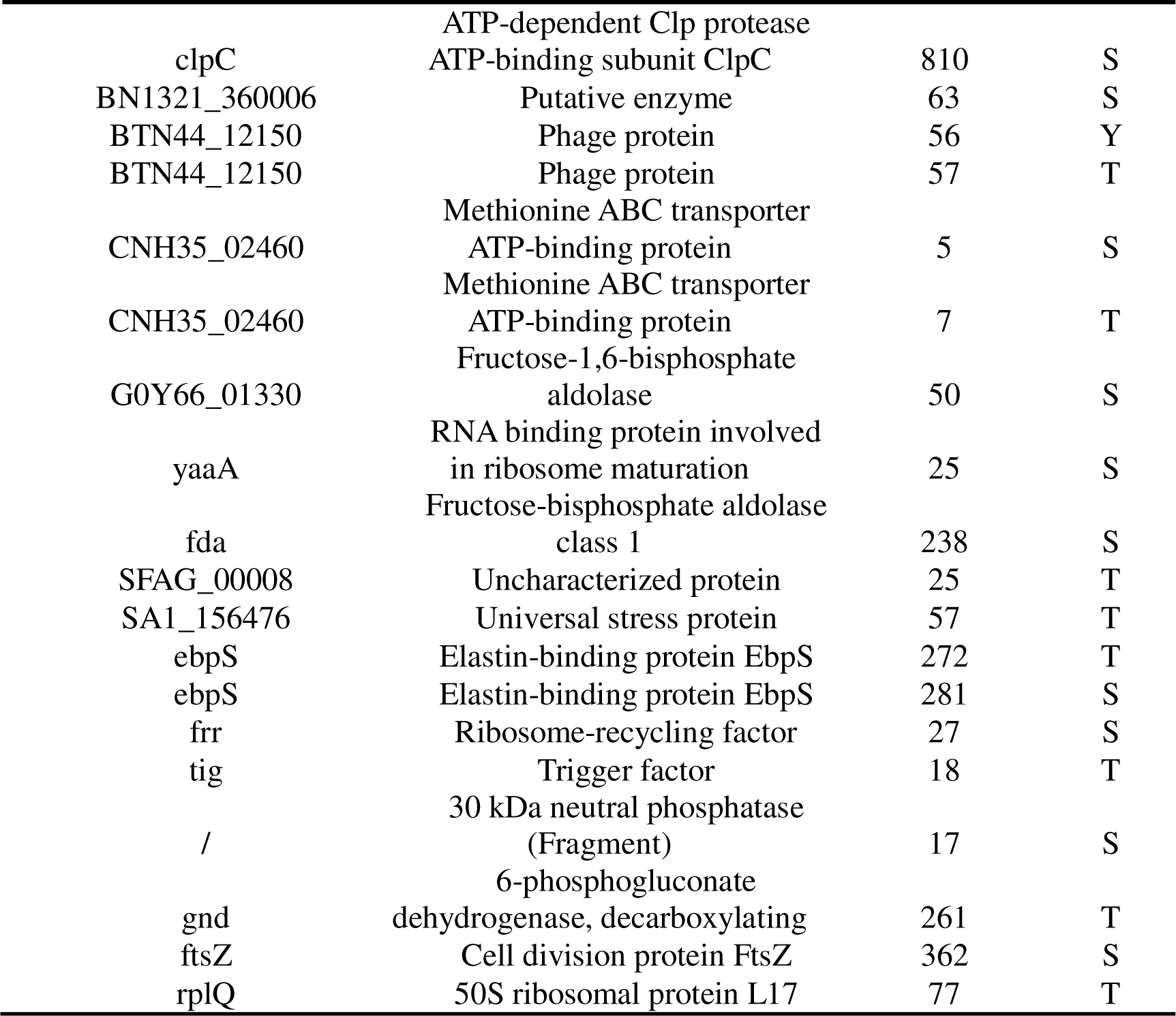
Differential proteins for sensitive and antifreeze strains after freezing (upregulated)

**Table S4.** Differential proteins for sensitive and antifreeze strains after freezing (downregulated)

| 基因名称 | 蛋白名称 | 位点 | 氨基酸 |
| --- | --- | --- | --- |
| V070_01283 | Uncharacterized protein | 758 | T |
| V070_01283 | Uncharacterized protein | 762 | T |
| V070_01283 | Uncharacterized protein | 746 | S |
| V070_01283 | Uncharacterized protein | 751 | S |
| V070_01283 | Uncharacterized protein | 763 | T |
| V070_01283 | Uncharacterized protein | 791 | S |
| V070_00528 | CBS domain-containing protein | 4 | T |
| V070_00528 | CBS domain-containing protein | 2 | Y |
|  | Fructose-bisphosphate aldolase |  |  |
| fda | class 1 | 238 | S |
| HMPREF0776_04 | Pyridoxamine 5'-phosphate |  |  |
| 35 | oxidase family protein | 4 | S |
|  | D-fructose-6-phosphate |  |  |
| GO788_00055 | amidotransferase | 5 | Y |
| GO788_00055 | D-fructose-6-phosphate<br>amidotransferase | 6 | T |
| GO788_00055 | D-fructose-6-phosphate<br>amidotransferase | 12 | T |
| G0V76_02915 | Uncharacterized protein | 151 | S |
| G0V76_02915 | Uncharacterized protein | 157 | Y |
| G0V76_02915 | Uncharacterized protein | 156 | S |
| pflB | Formate acetyltransferase | 650 | S |
| SFAG_02032 | DNA-binding response regulator | 210 | S |
| SFAG_02032 | DNA-binding response regulator | 214 | T |
| CV021_12290 | DUF1828 domain-containing<br>protein (Fragment) | 8 | S |
| CV021_12290 | DUF1828 domain-containing<br>protein (Fragment) | 5 | T |
| gluD | NAD-specific glutamate<br>dehydrogenase | 267 | T |

**Table S5.**
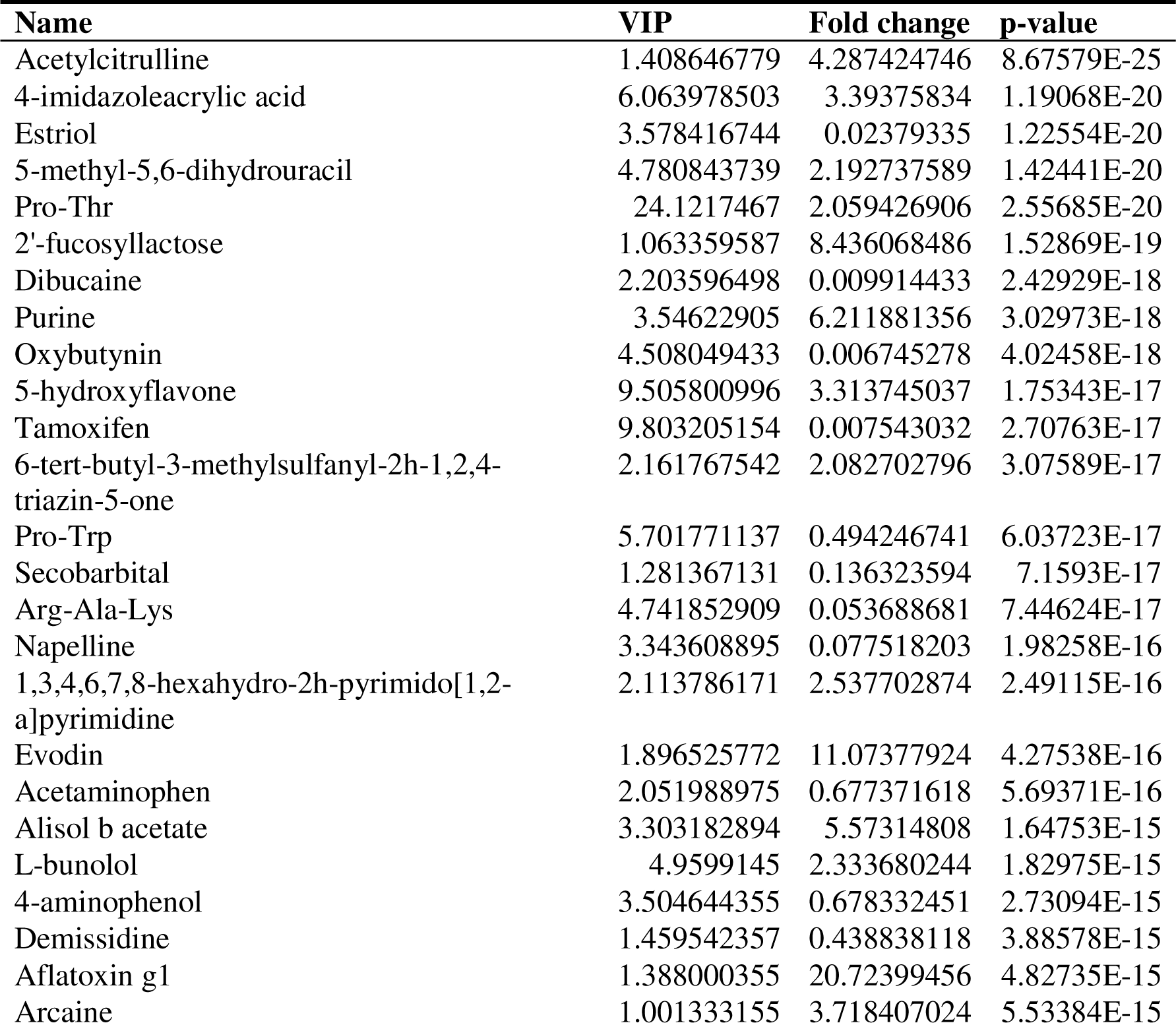
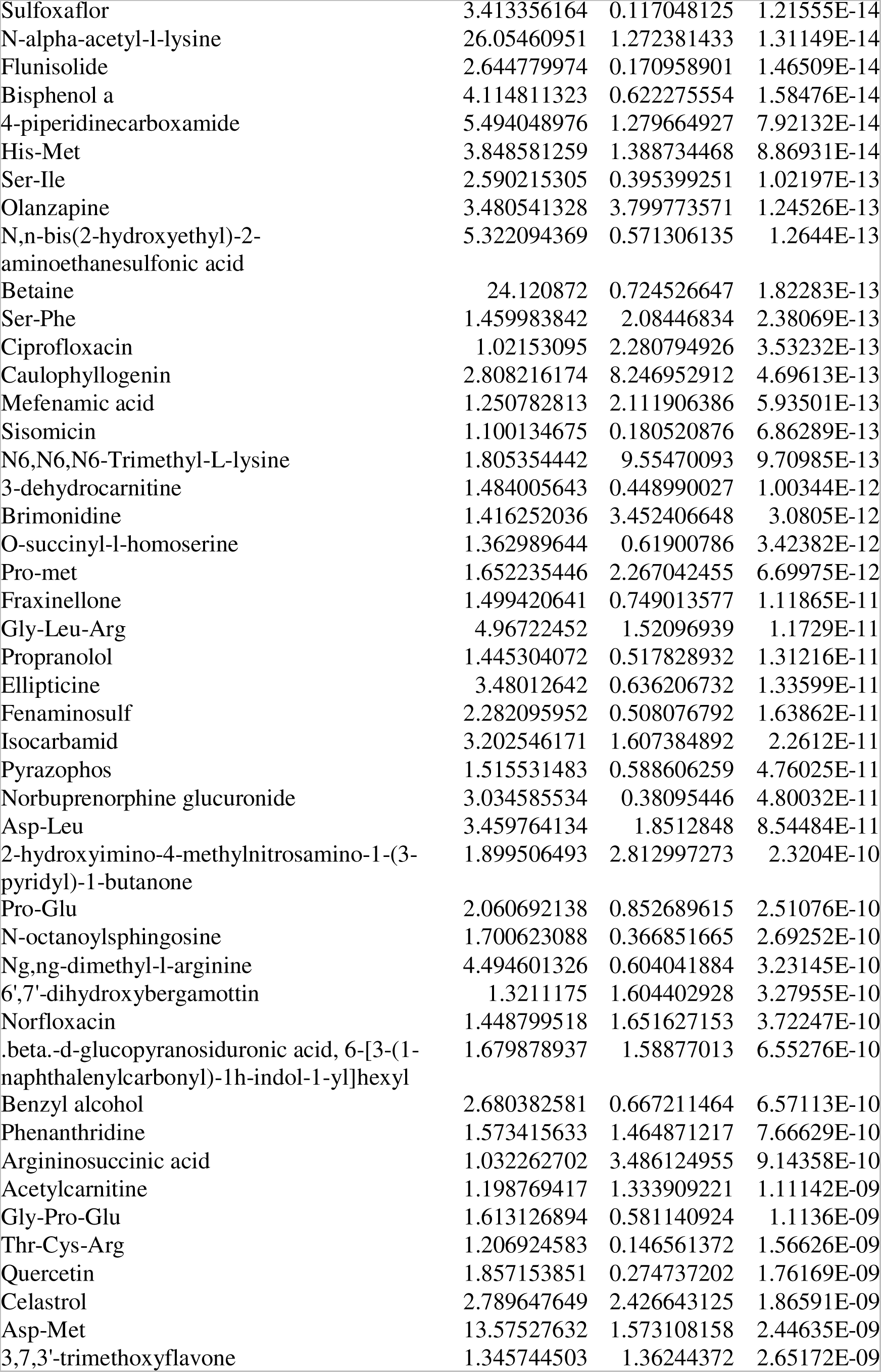
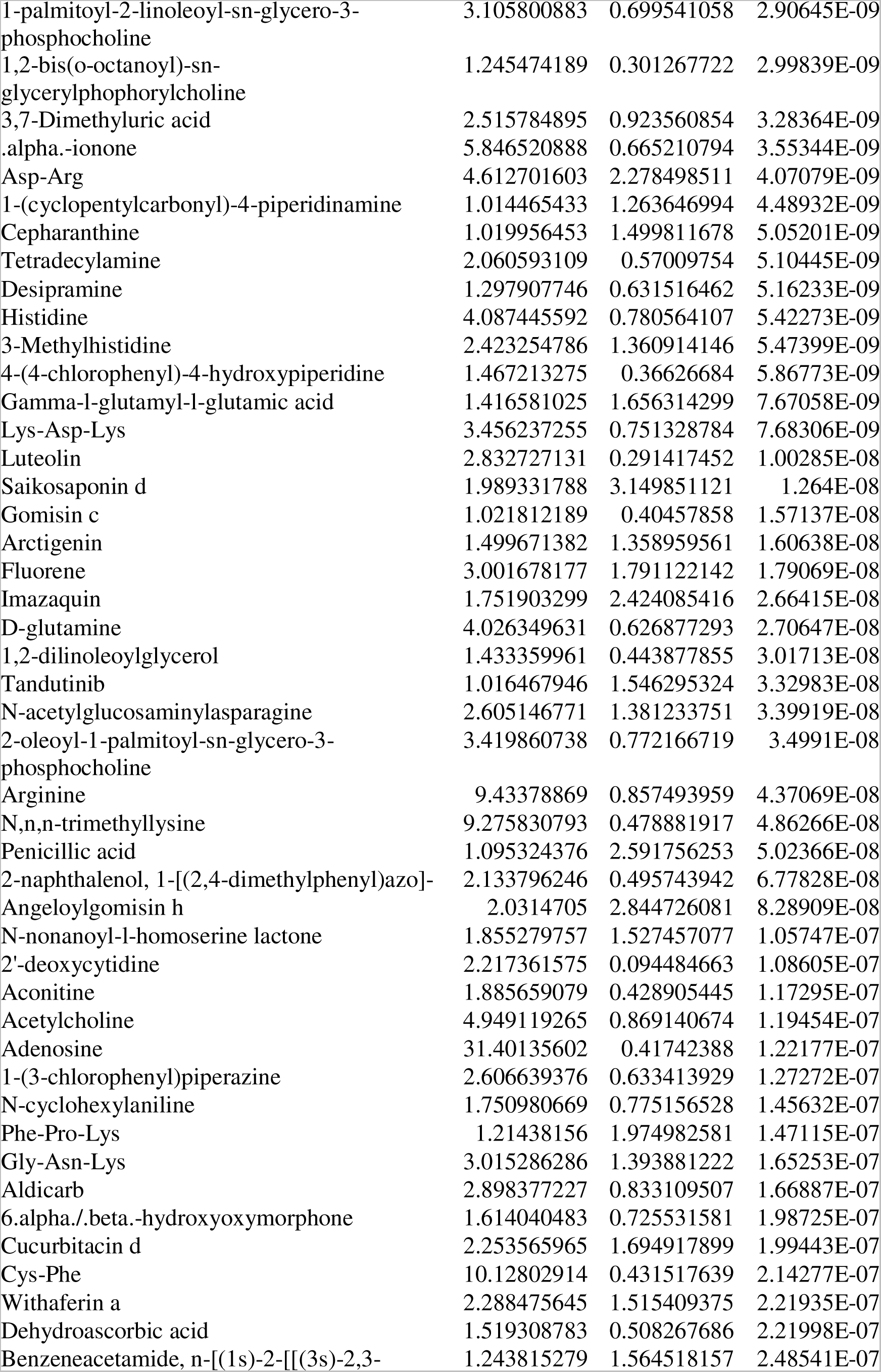
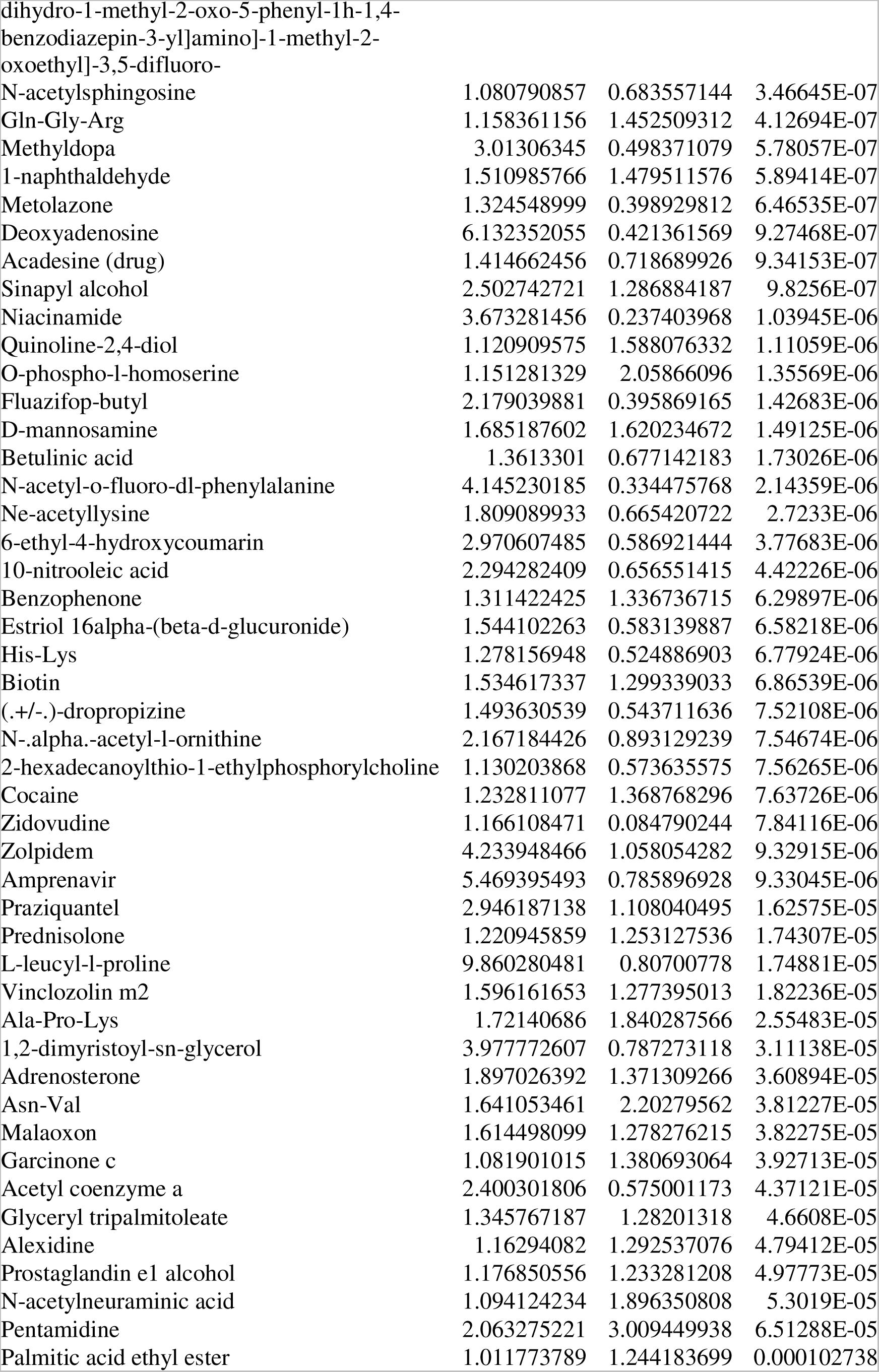
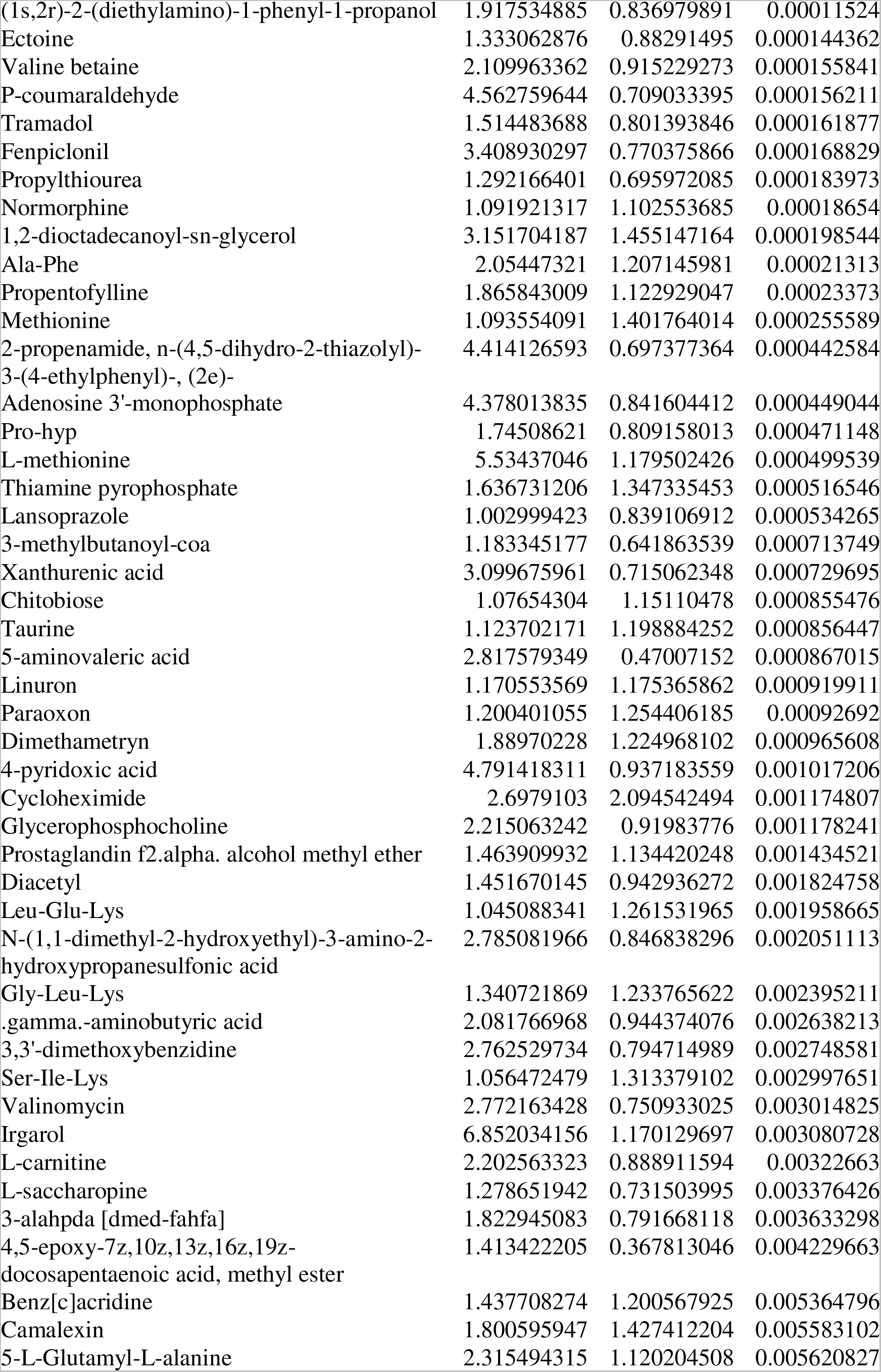
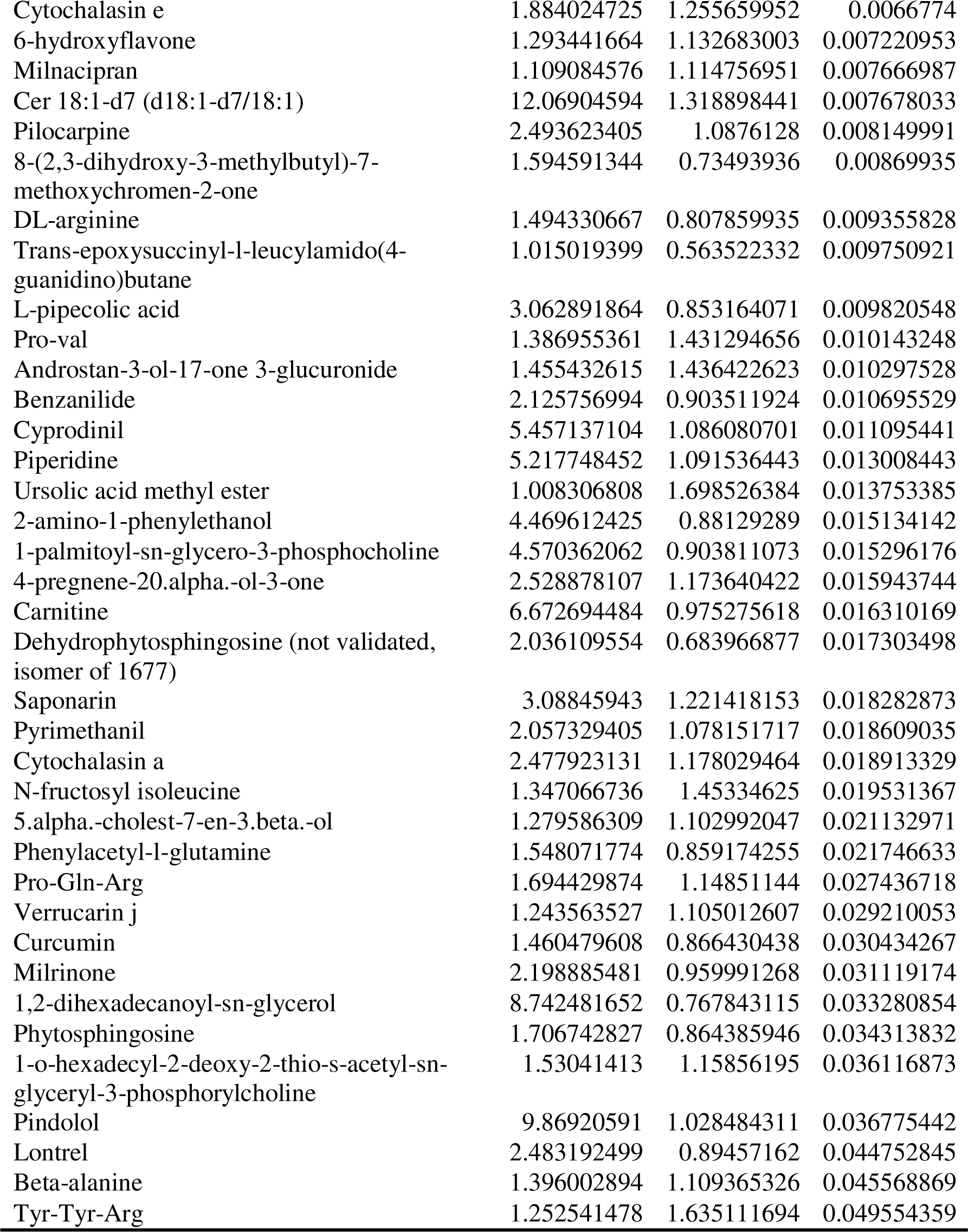
Staphylococcus aureus H7 vs H8 differential metabolites before freezing in positive ion mode.

**Table S6.**
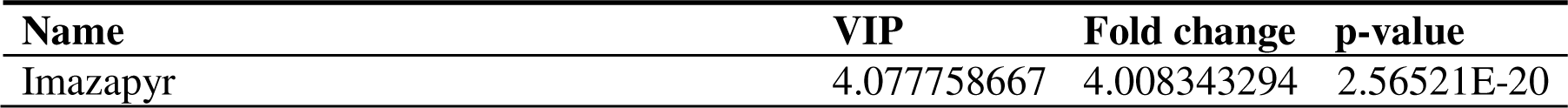
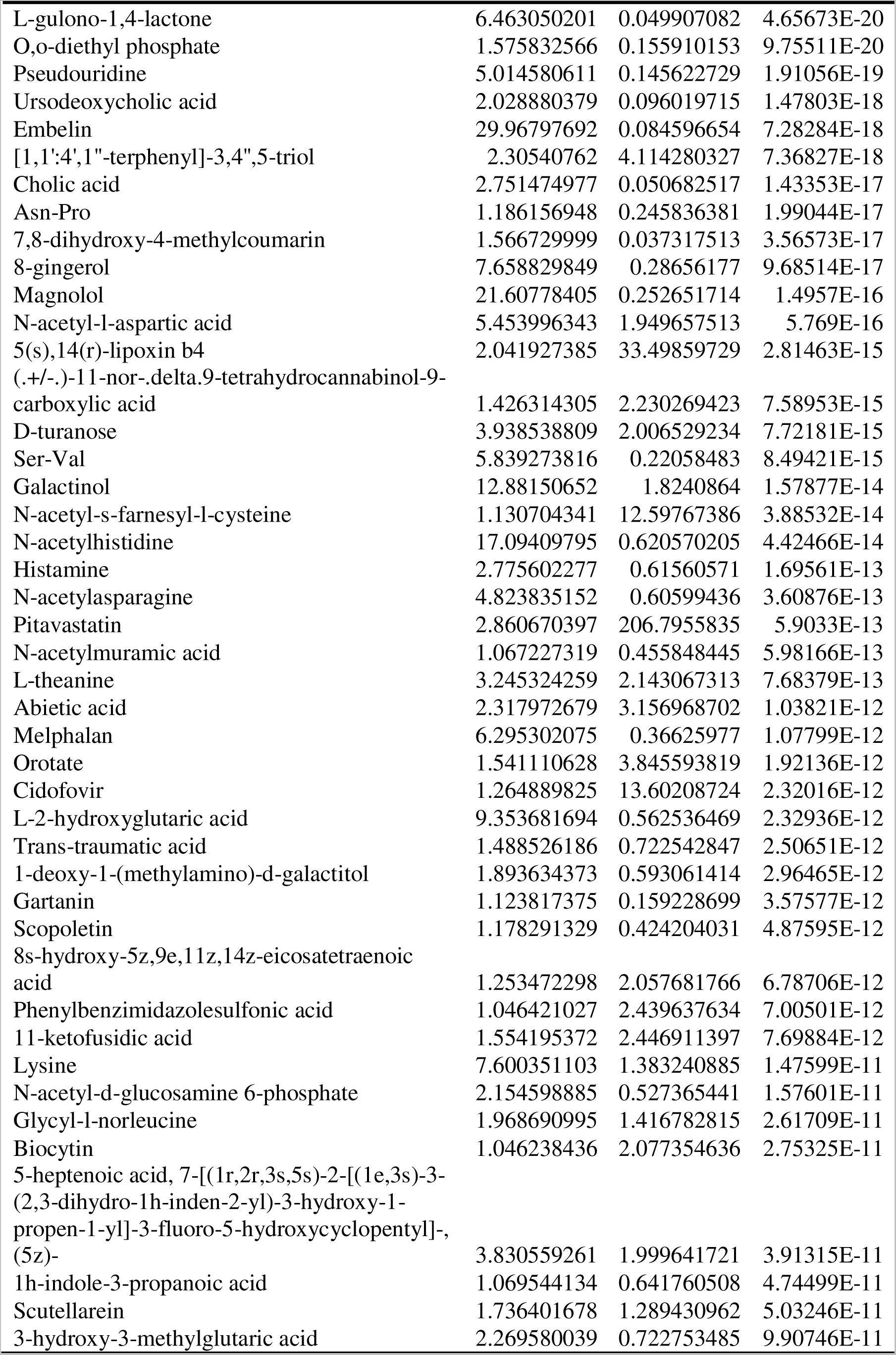
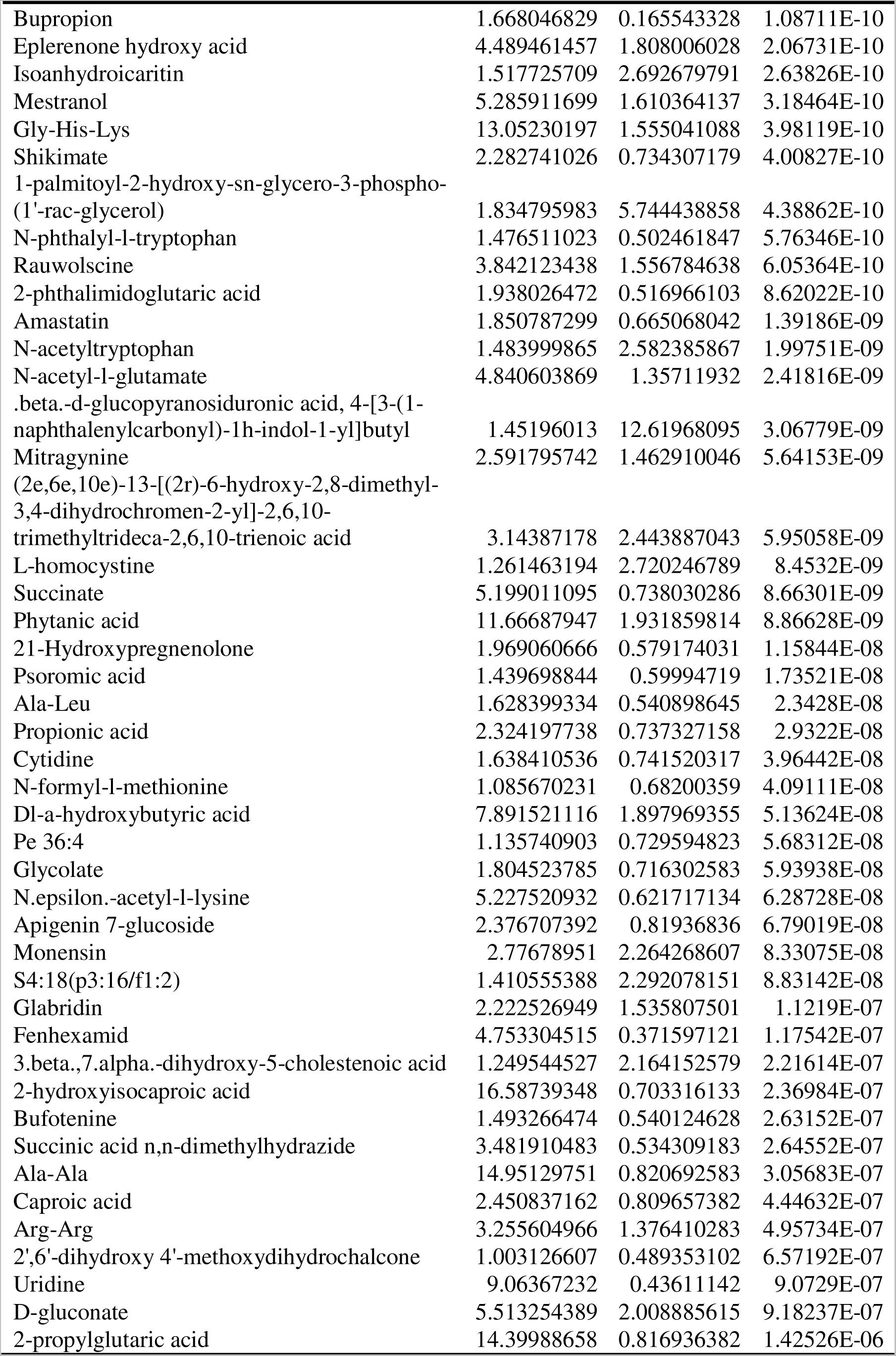
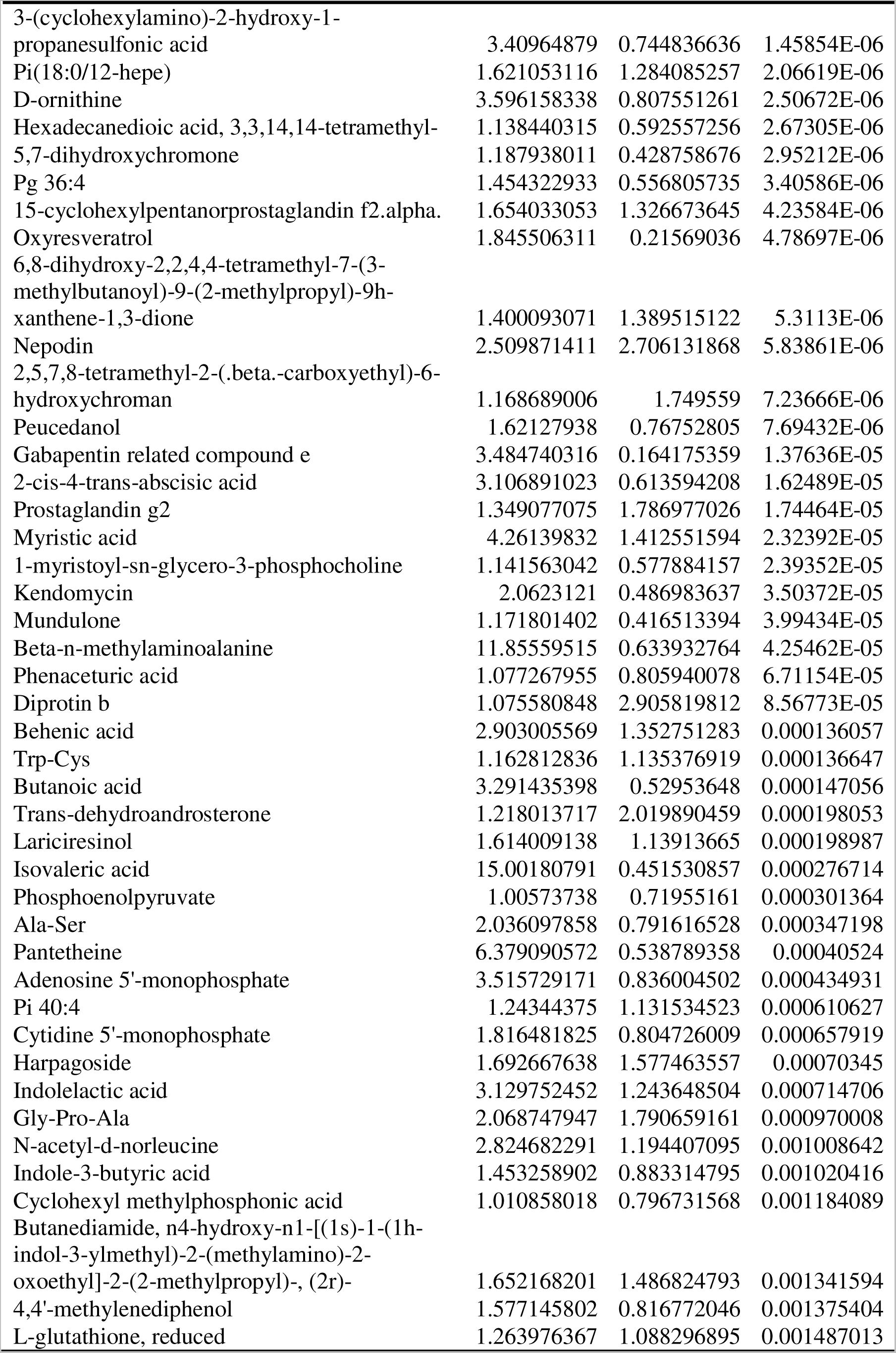
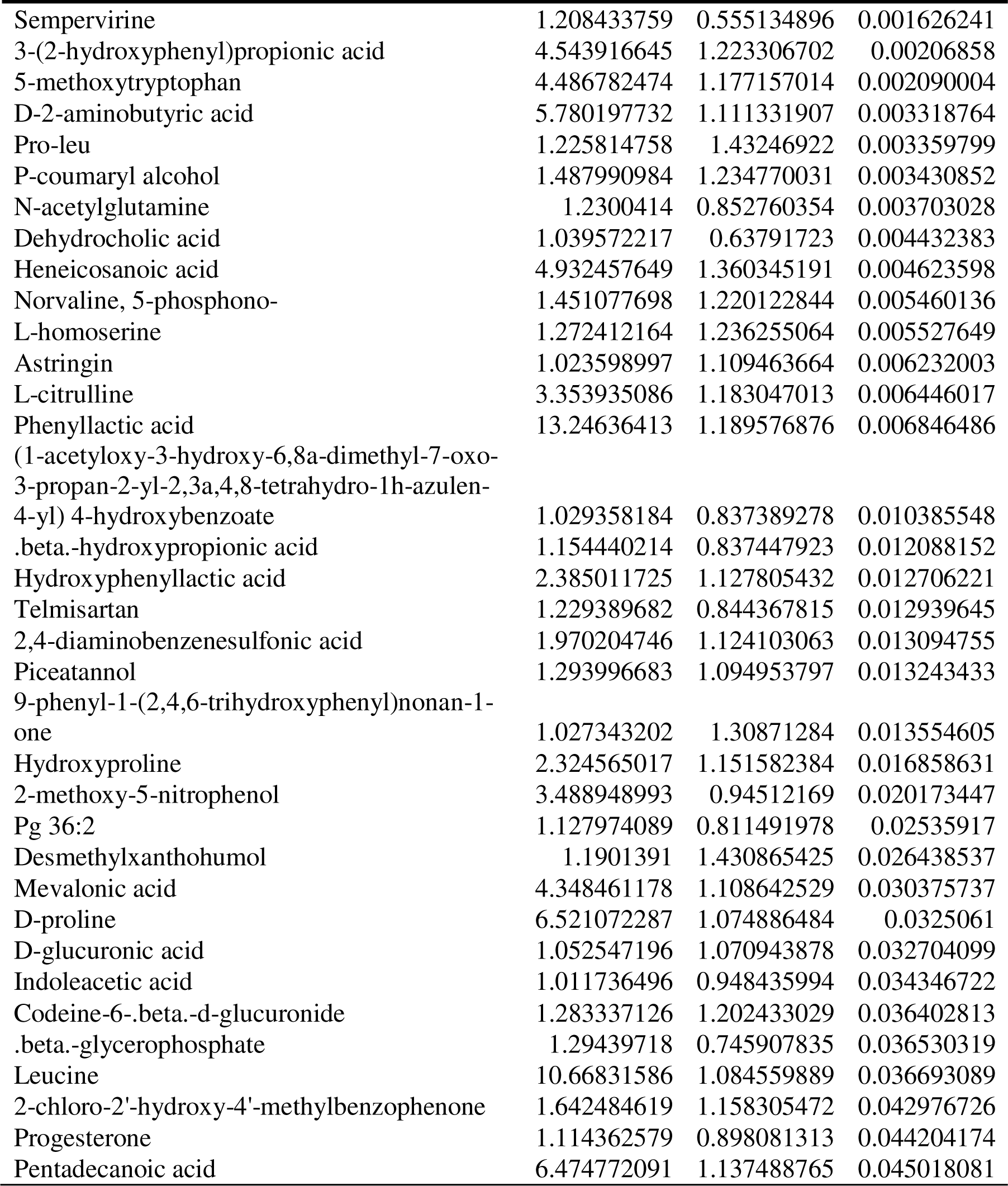
Staphylococcus aureus H7 vs H8 differential metabolites before freezing in negative ion mode.

**Table S7.**
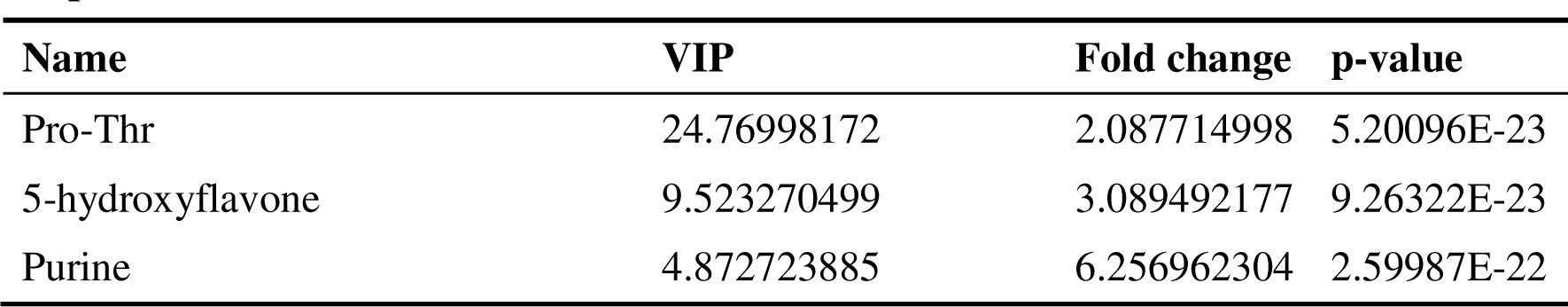
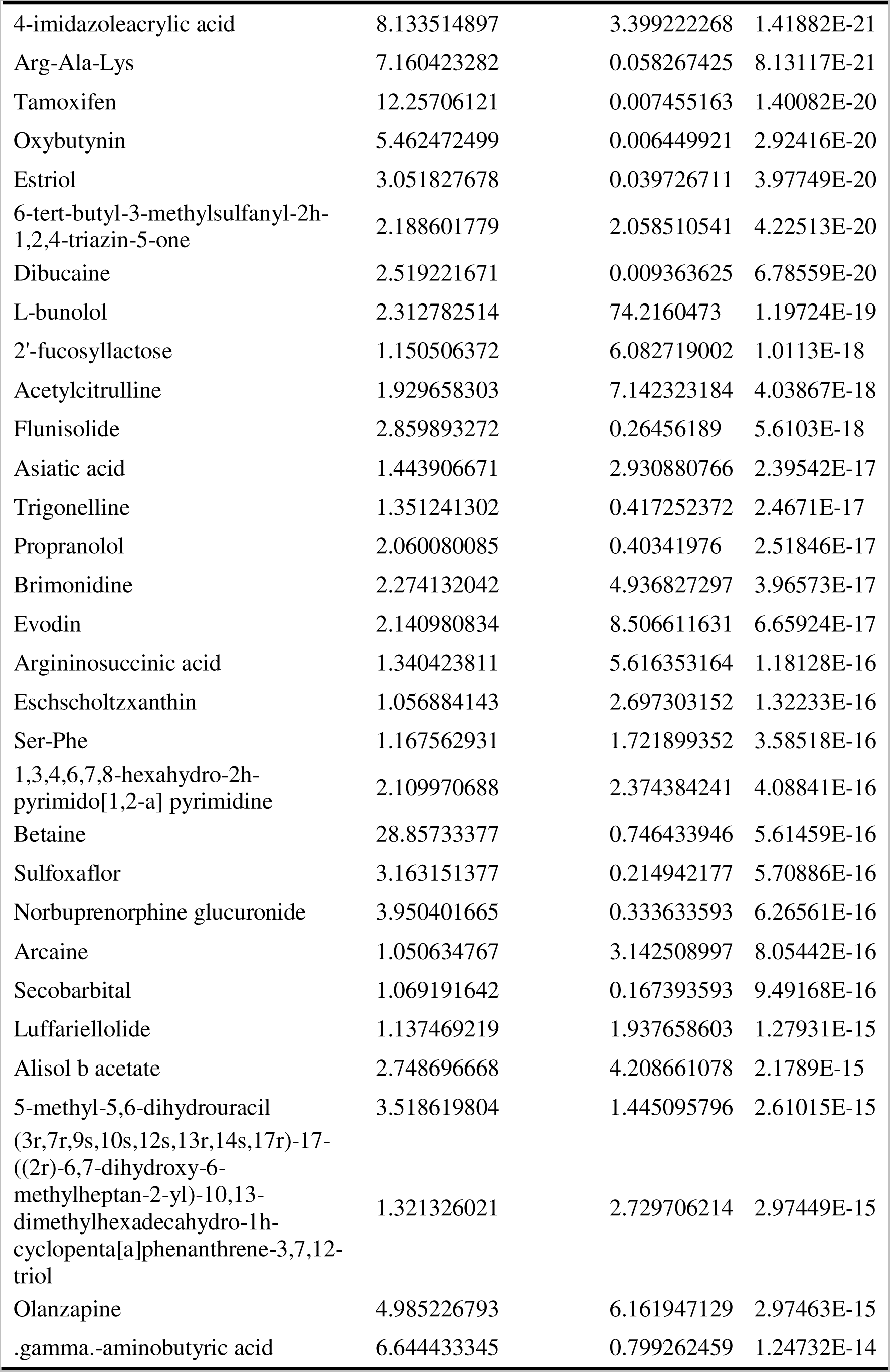
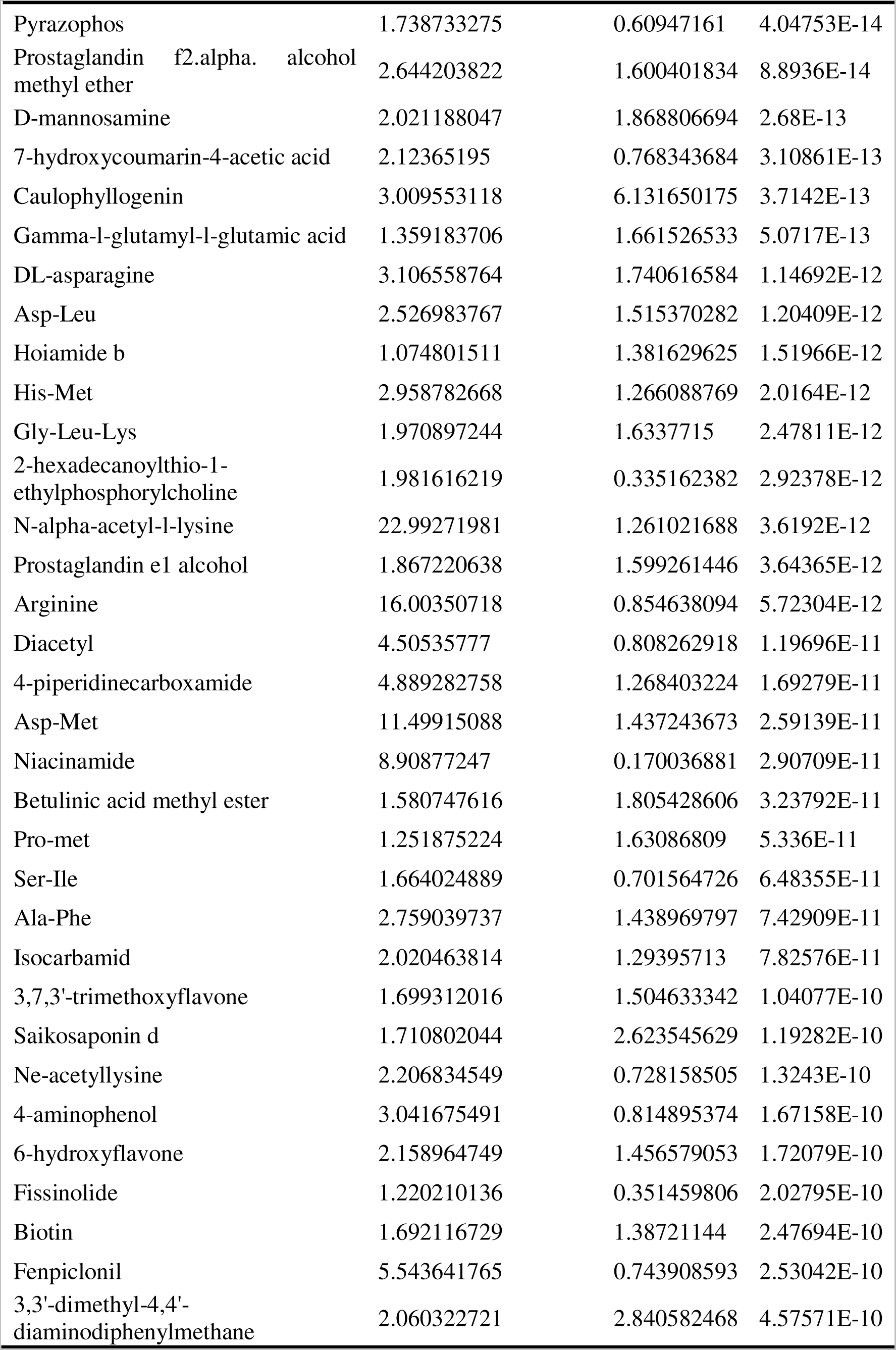
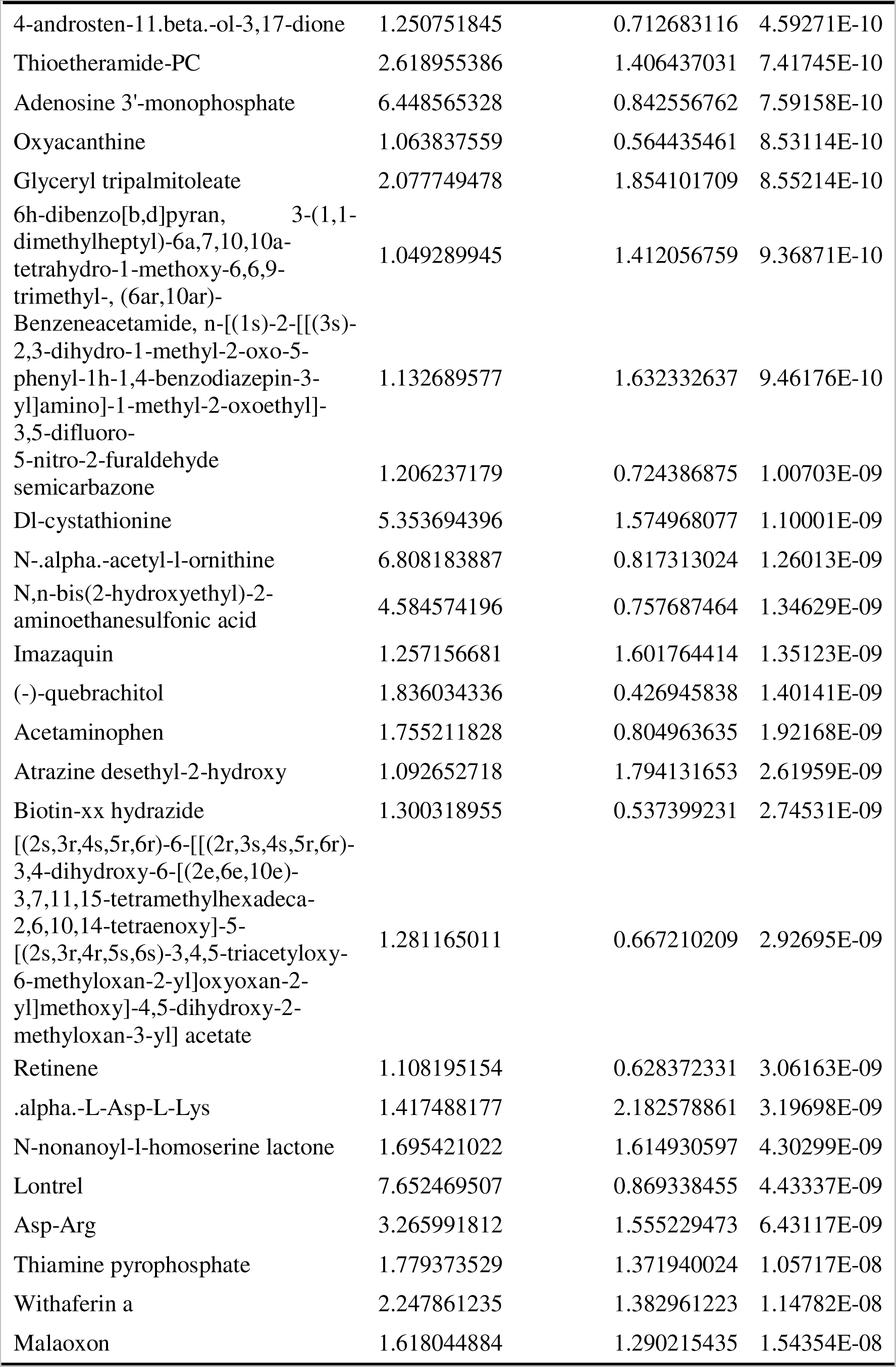
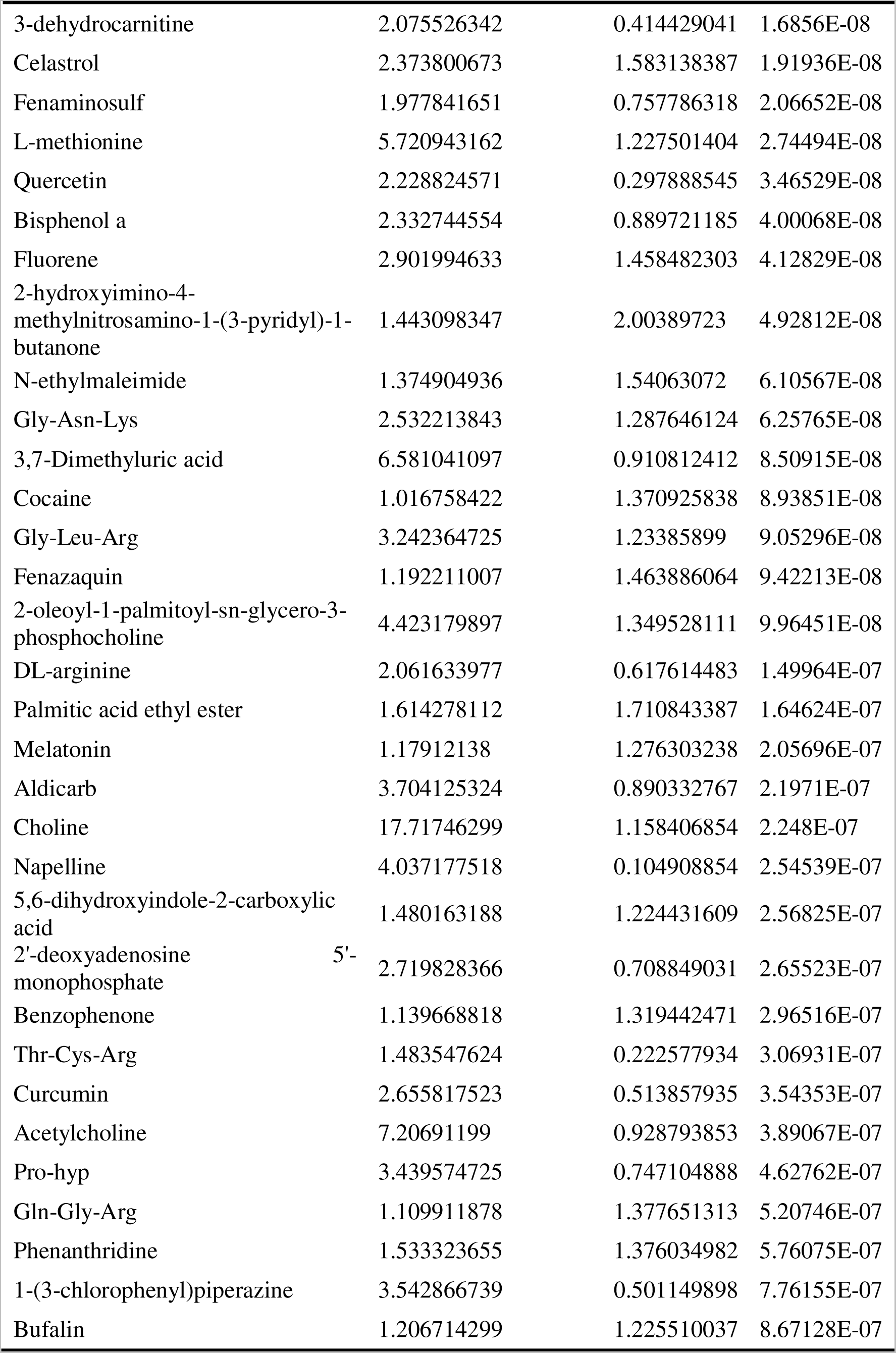
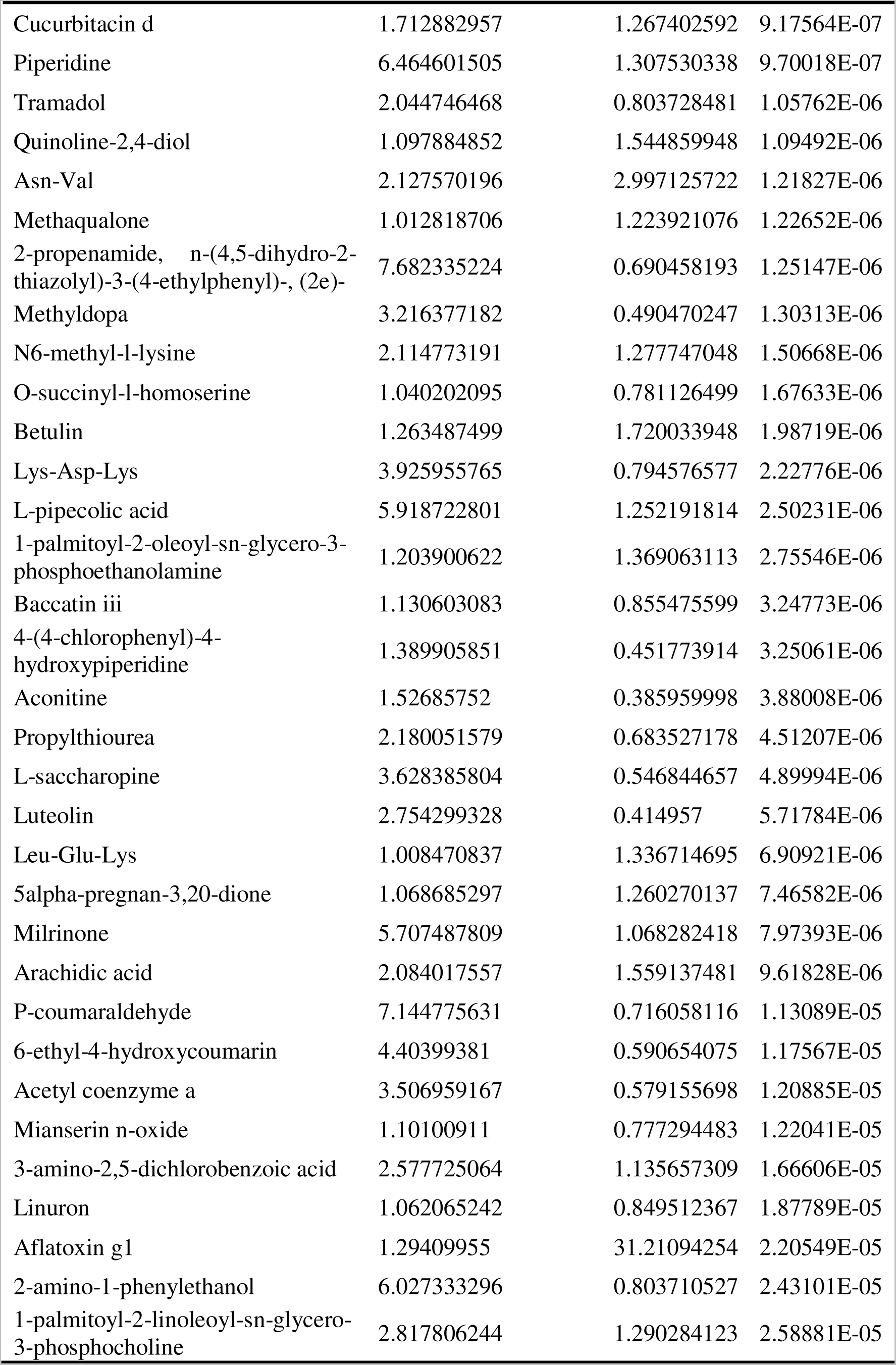
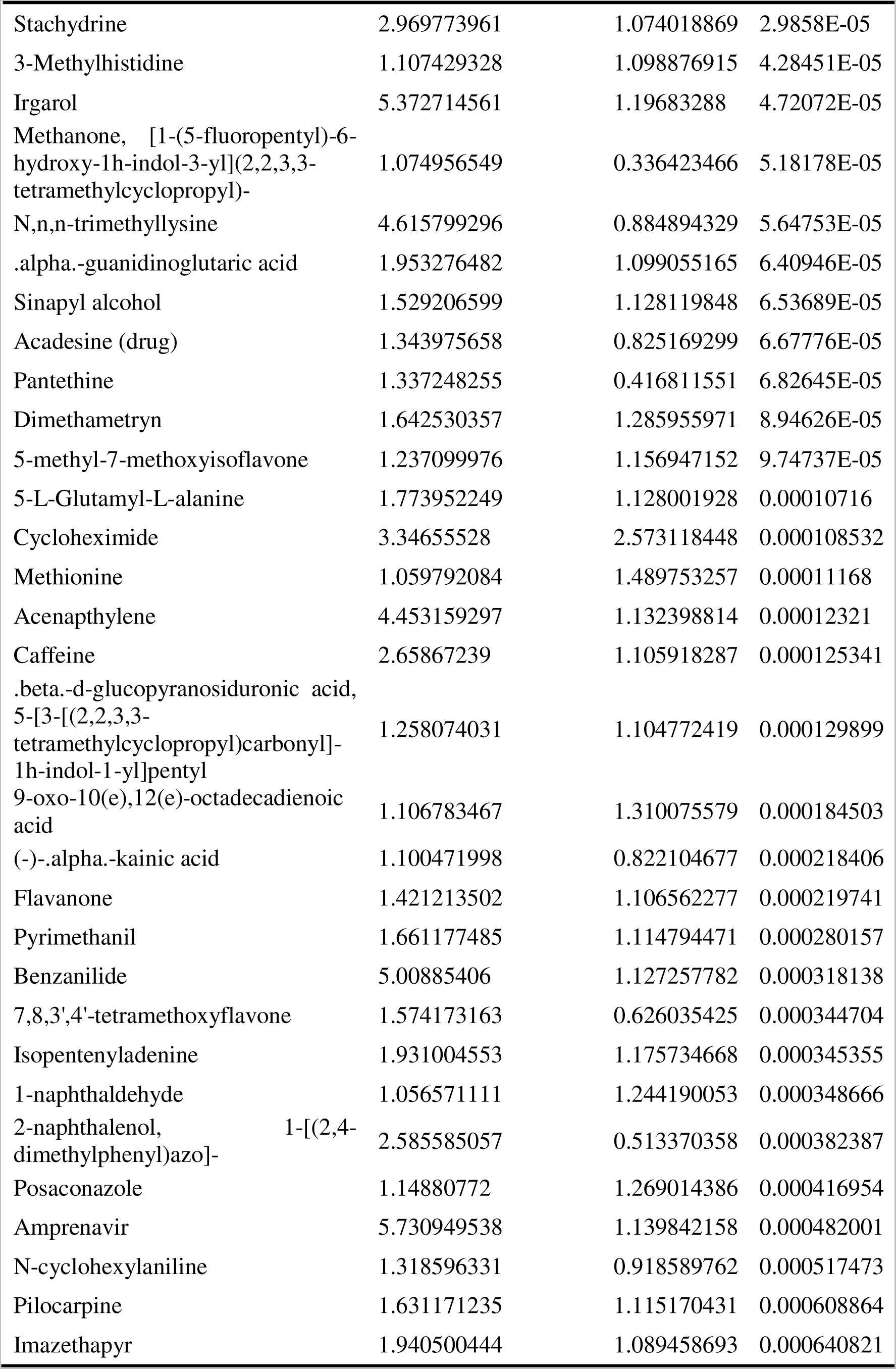
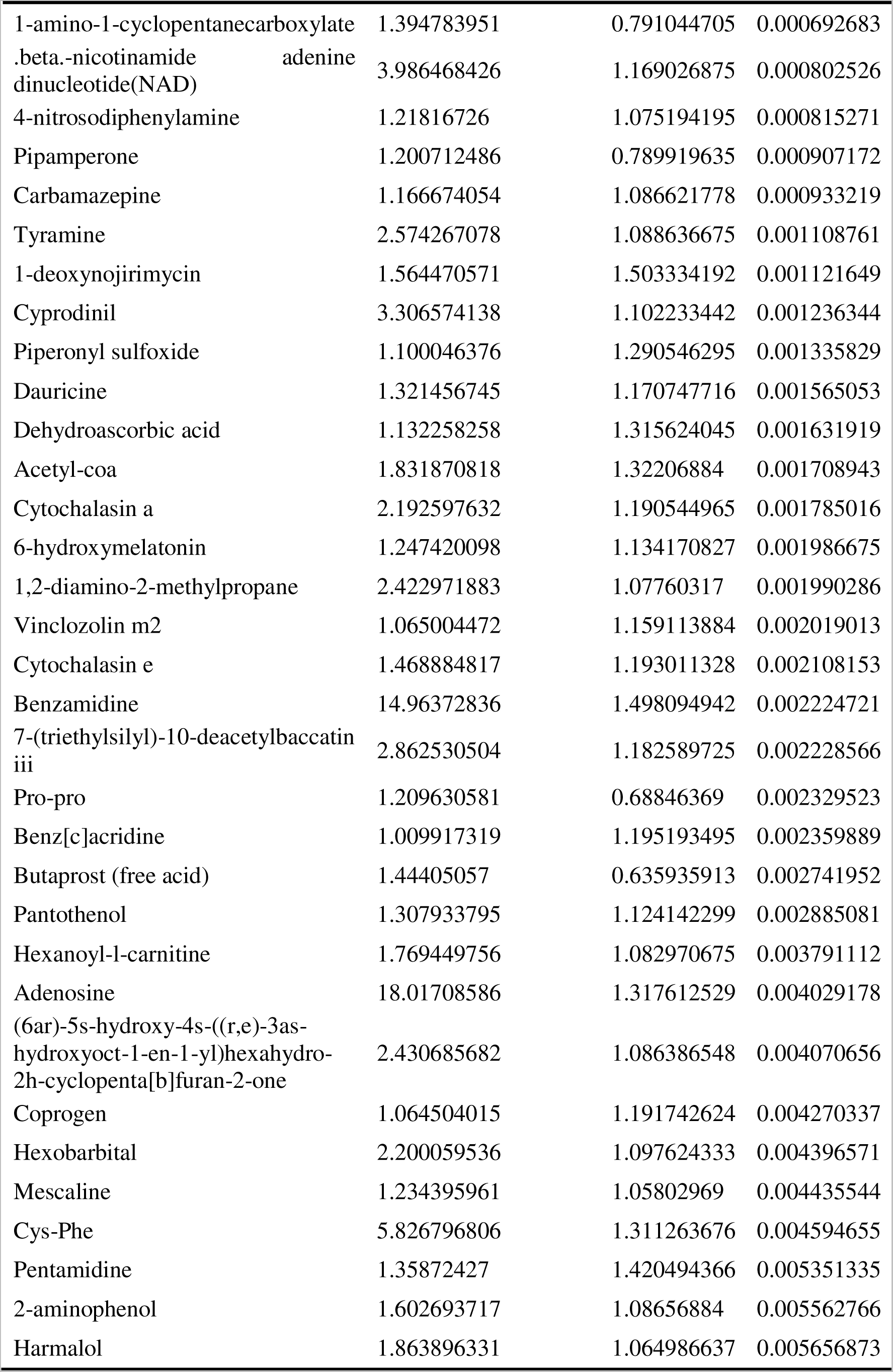
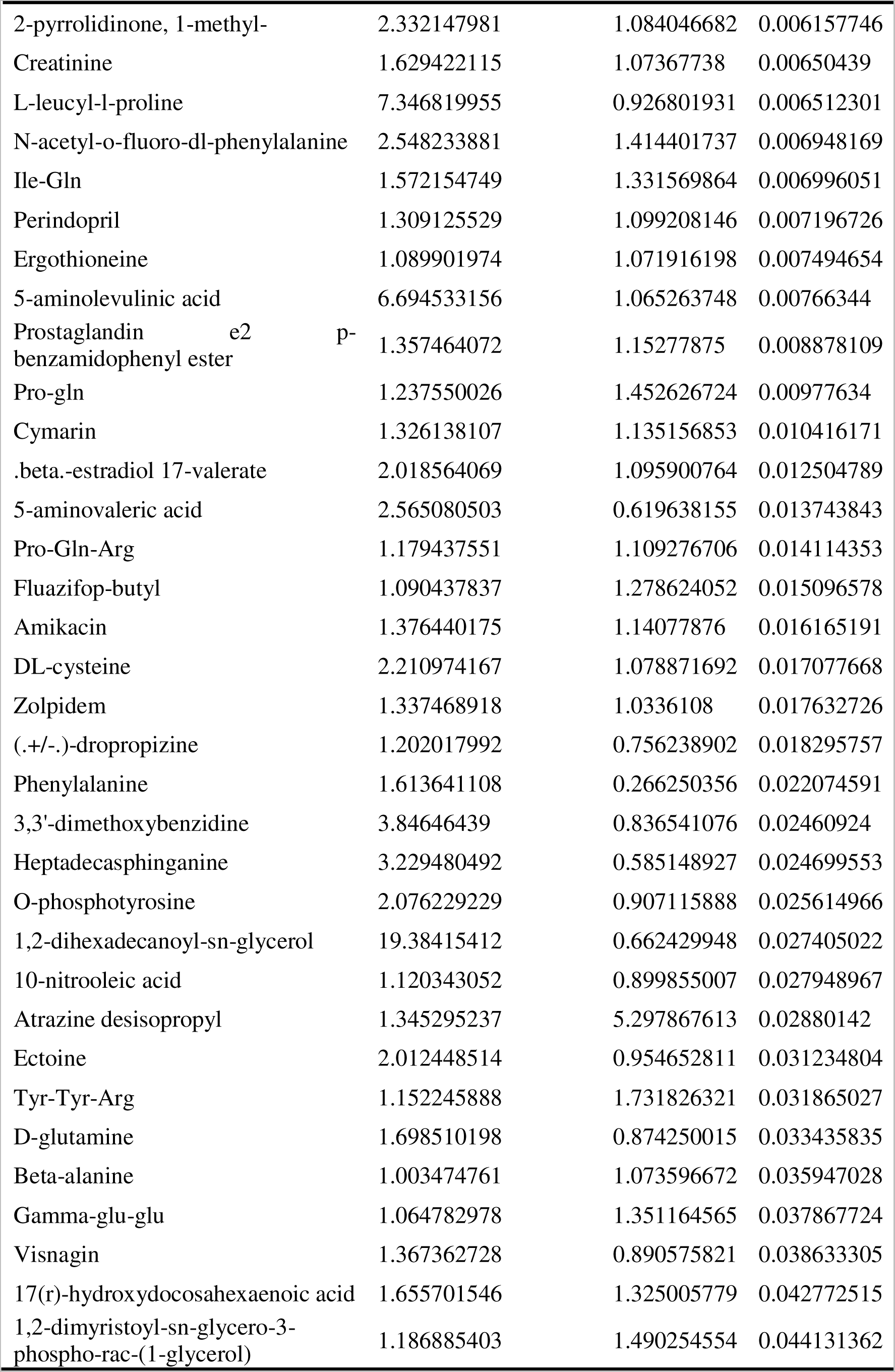
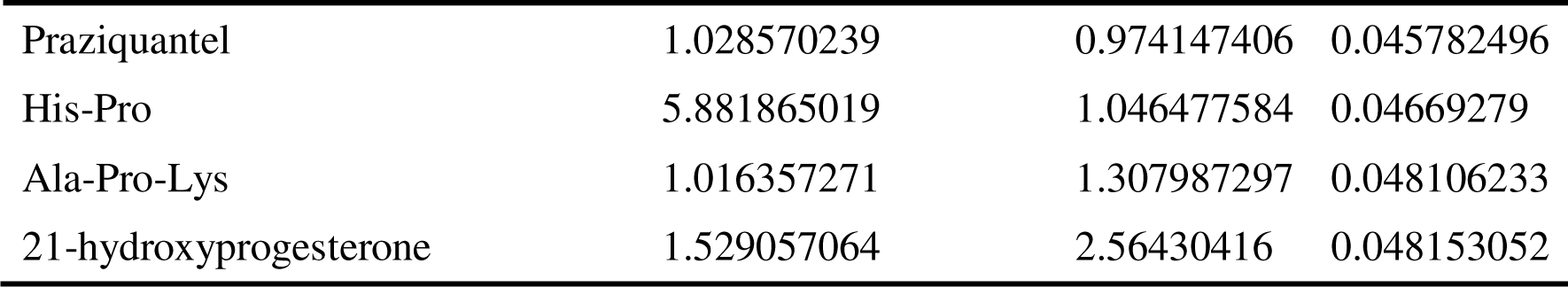
*Staphylococcus aureus* Q7 vs Q8 differential metabolites after freezing in positive ion mode.

**Table S8.**
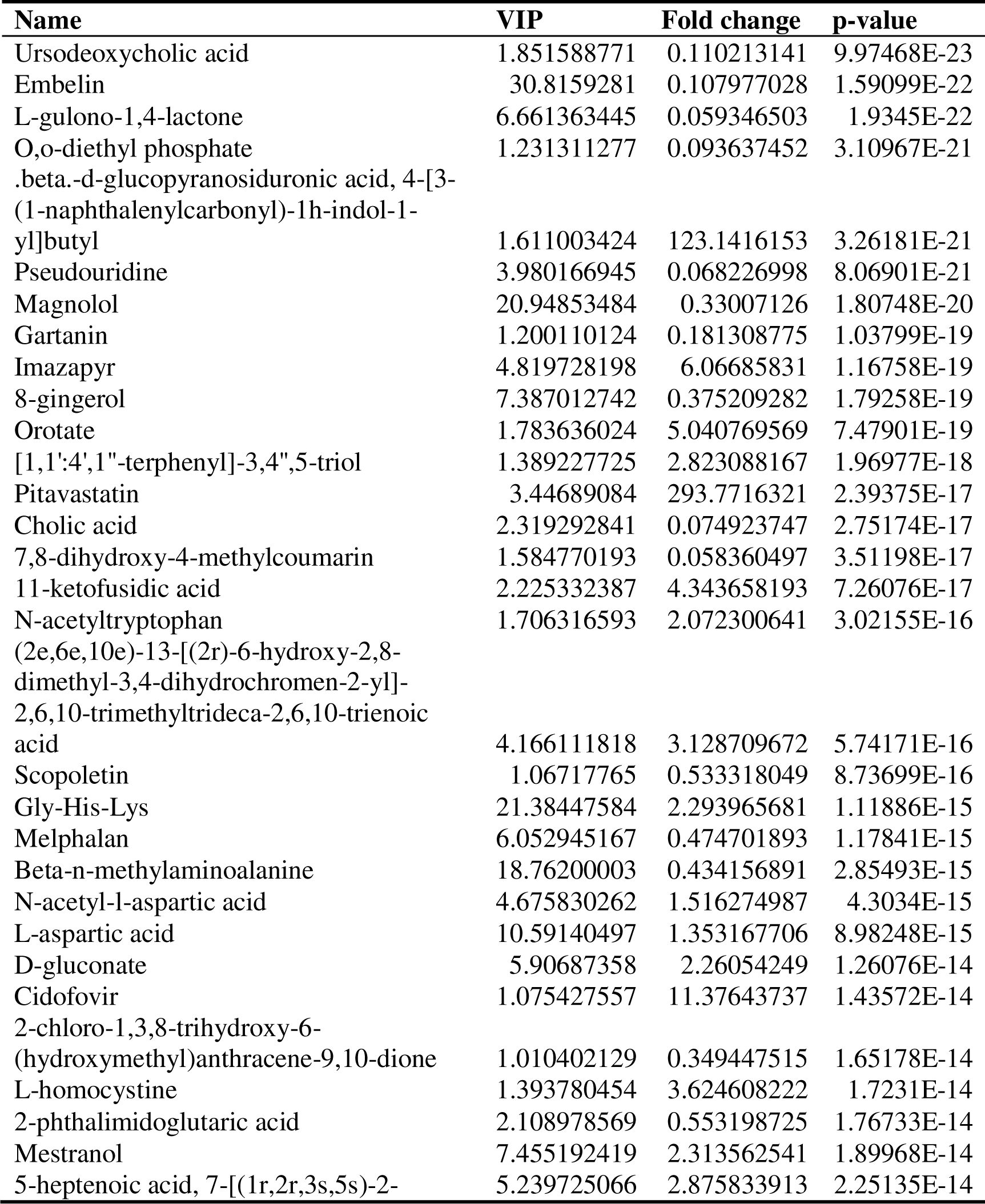
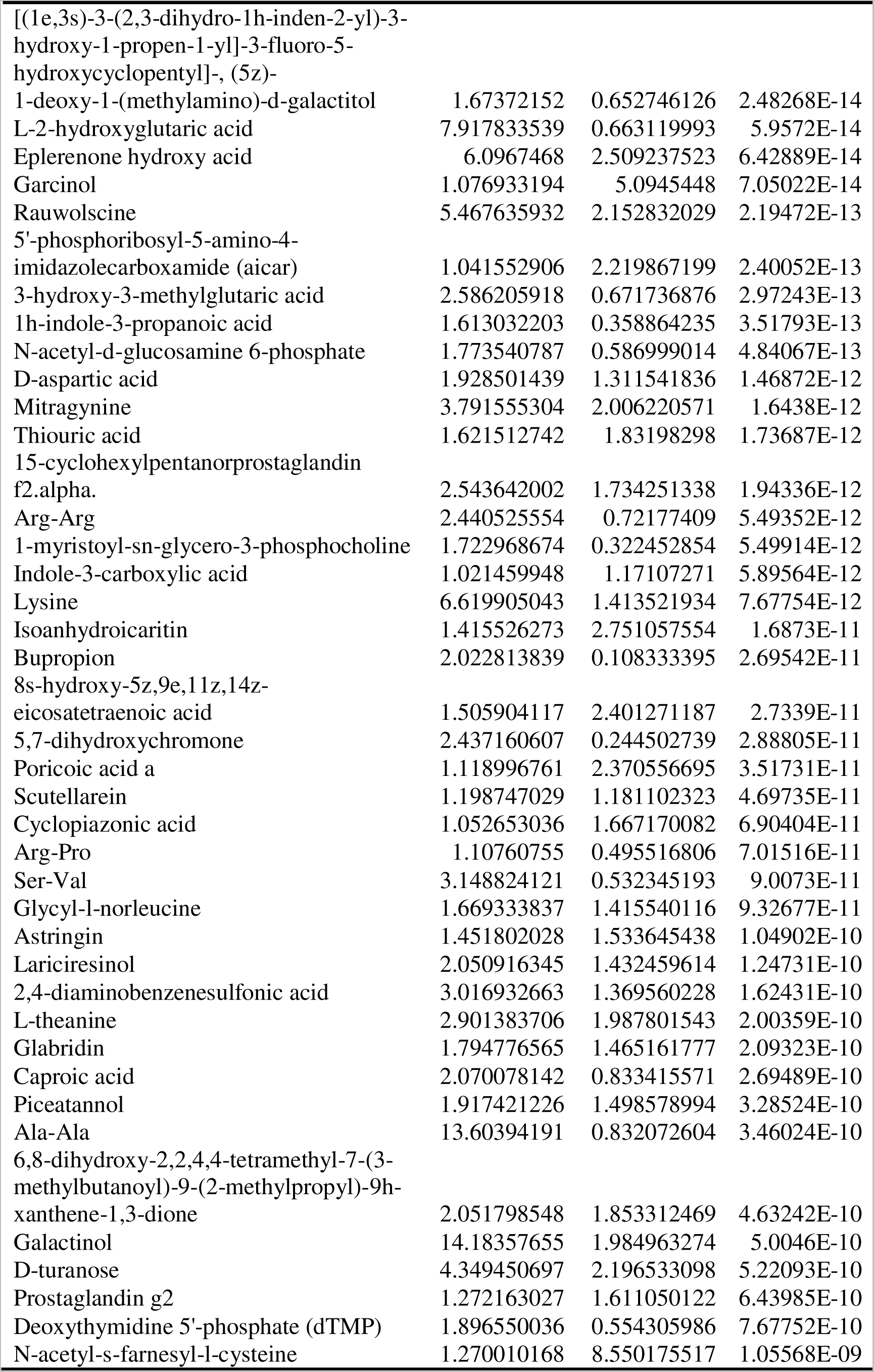
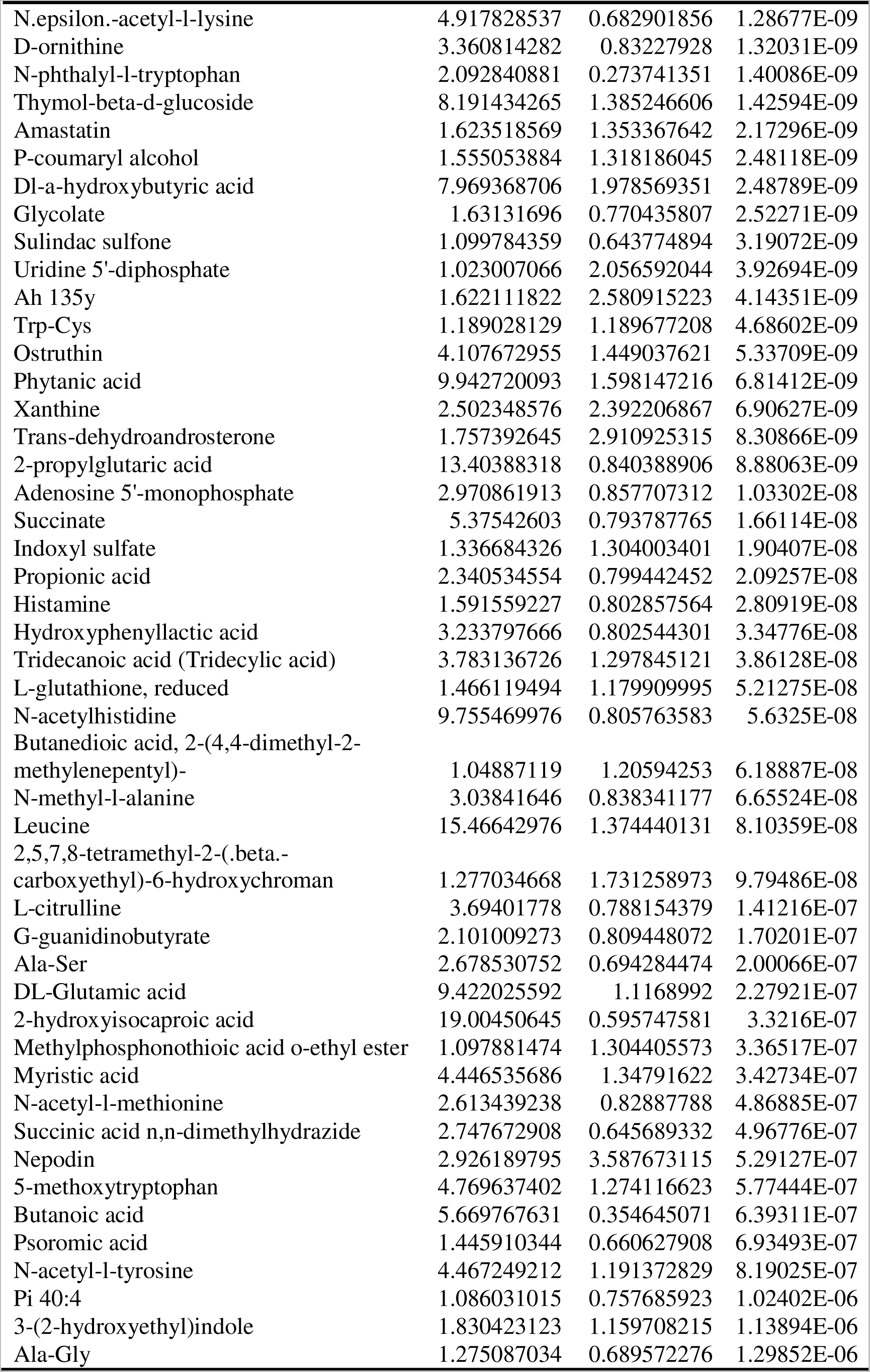
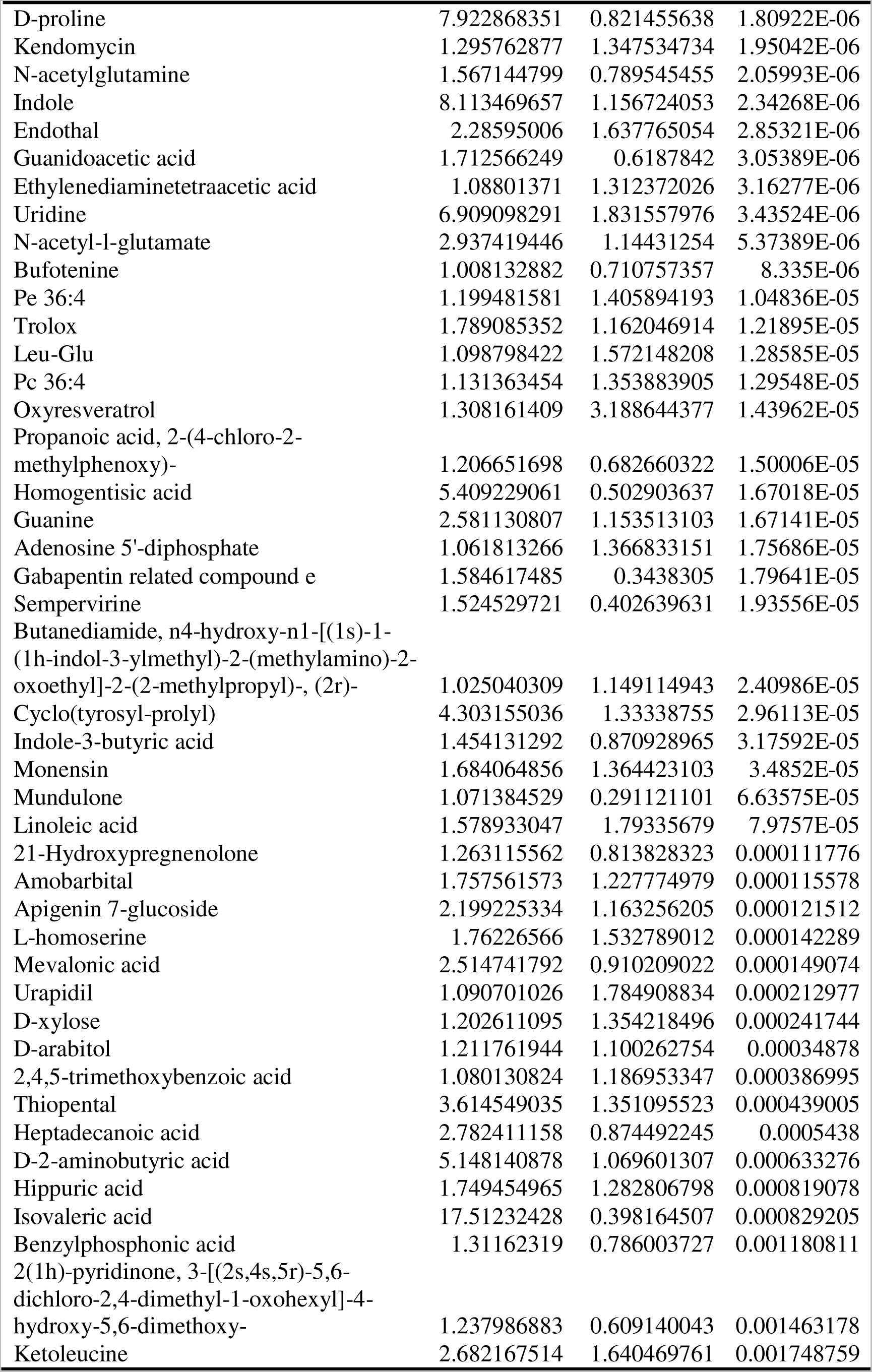
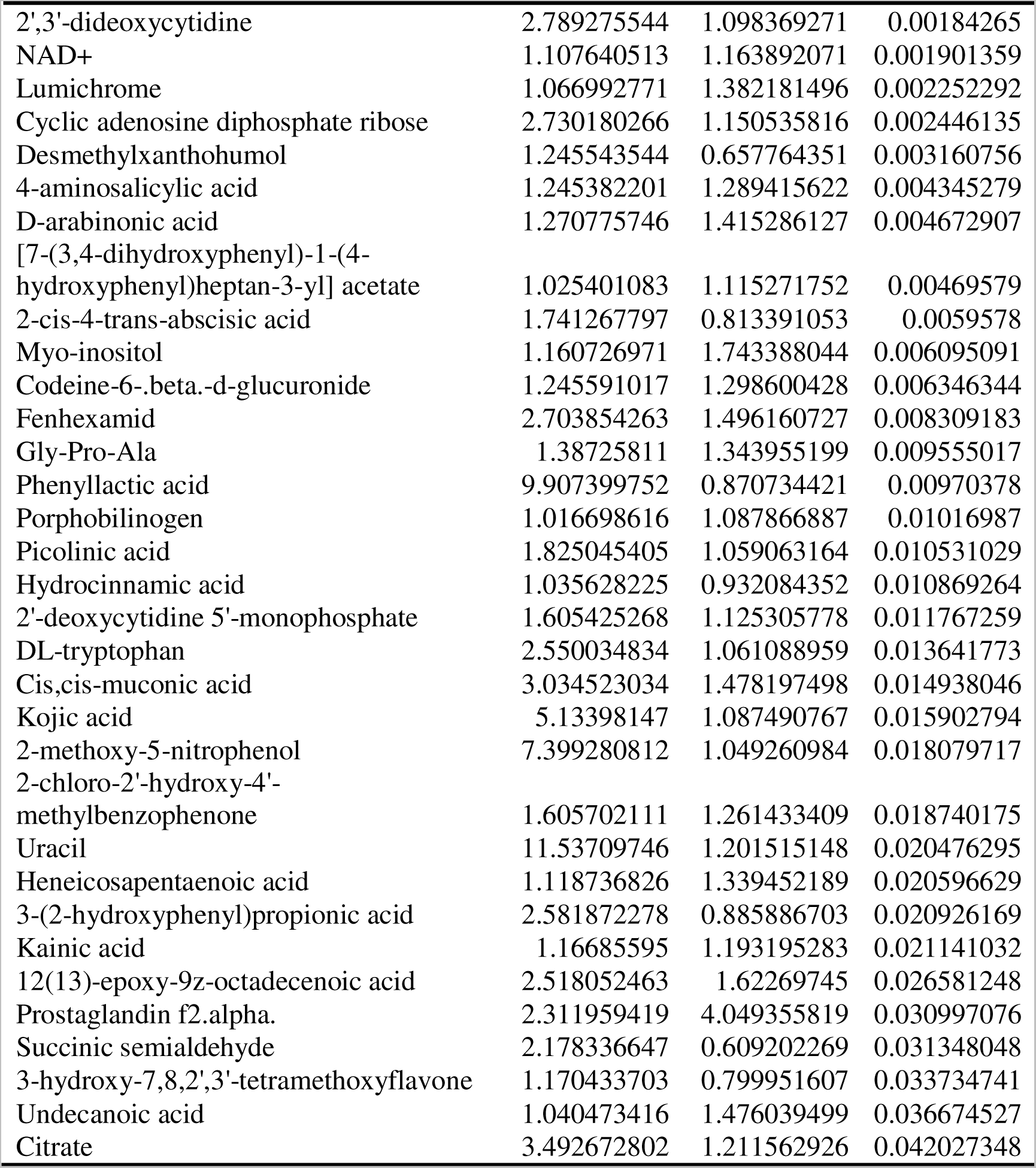
Staphylococcus aureus Q7 vs Q8 differential metabolites after freezing in nagetive ion mode.

